# ANFIS-based fuzzy systems for searching dna-protein binding sites

**DOI:** 10.1101/058800

**Authors:** Dianhui Wang, Monther Alhamdoosh, Witold Pedrycz

## Abstract

Transcriptional regulation mainly controls how genes are expressed and how cells behave based on the transcription factor (TF) proteins that bind upstream of the transcription start sites (TSSs) of genes. These TF DNA binding sites (TFBSs) are usually short (5-15 base pairs) and degenerate (some positions can have multiple possible alternatives). Traditionally, computational methods scan DNA sequences using the position weight matrix (PWM) of a given TF, calculate binding scores for each K-mer against the PWM, and finally classify a K-mer as to whether it is a putative TFBS or a background sequence based on a cut-off threshold. The FSCAN system, which is proposed in this paper, employs machine learning techniques to build a learner model that is able to identify TFBSs in a set of bound sequences without the need for a cut-off threshold. Our proposed method utilizes fuzzy inference techniques along with a distribution-based filtering algorithm to predict the binding sites of a TF given its PWM model and phastCons scores for the input DNA sequences. Data imbalance reduction techniques are also used to ease the learning of the adaptive-neuro fuzzy inference system (ANFIS) algorithm. The proposed system is tested on 22 ChIP-chip sequence-sets from the Saccharomyces Cerevisiae genome. Our results show that FSCAN outperforms other approaches like MatInspector and MATCH and is quite robust. As more transcriptional data becomes available, our proposed framework encourages the use of fuzzy logic techniques in the prediction of TFBSs.

## 1 Introduction

Transcription is a key stage in the gene expression process in which DNA nucleotides of genes are transliterated into RNA nucleotides by RNA polymerase. Some of these RNAs are then used to synthesize functional proteins while others operate in their RNA molecular state. These functional compounds determine the cell functionalities and morphology. However, the amount of protein or RNA that is produced from a particular gene is mainly regulated by specific proteins called transcription factors (TFs). Transcription factor proteins bind to short and degenerate DNA elements (5-15 base pairs) called transcription factor binding sites (TFBSs). TFBSs commonly occur in the promoter and upstream (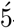-UTR) regions of genes in prokaryotic and eukaryotic organisms, e.g., yeast. Nevertheless, some TFs may also bind to 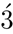-UTR or intronic regions in high eukaryotes, e.g., human genome. Knowledge of the exact locations of transcription factor binding sites enables researchers to better understand the gene regulatory networks of many organisms and consequently design drugs for many diseases, especially cancers. Due to the shortness and degeneracy of TFBS sequences, the prediction of their exact locations *in silico* is a very challenging task yet it is very desirable. However, several in *vivo* techniques were proposed to study the DNA-protein interactions in whole genomes. Most of these technologies are based on DNA microarrays. For example, chromatin immunoprecipitation on chip (ChIP-chip) Ren *et al.* (2000) is the most widely used method for identifying TFBSs although it lacks the ability to determine TF binding specificity effectively because of the high signal-to-noise ratio. Another approach is protein binding microarray (PBM) Berger and Bulyk (2006); Bulyk (2007) which runs highly parallel microarray technology which enables it to characterize the DNA binding specificities of TFs in a very high-throughput manner. Unlike microarray-based methods, ChIP-Seq Johnson *et al.* (2007) employs massively parallel DNA sequencing technology in addition to chromatin immunoprecipitation in order to identify the locations of TFBSs at higher resolution. ChIP-based approaches usually use longer probe sequences than the PBM method.

The recent advancement in genome sequencing technologies has resulted in a vast amount of genome sequences but with limited knowledge of the gene regulatory networks of these genomes. The limitations of the aforementioned experimental approaches result in an urgent need to develop computational approaches that could help scientists in studying the TF protein-DNA interactions, i.e., locating putative transcription factor binding sites in genomes. However, developing such a computational system is not straightforward. Three key issues must be considered in order to build a reliable and precise system:(i) finding a good model for the DNA binding specificity of a transcription factor protein, (ii) modelling/scoring the binding affinity of a transcription factor to a given DNA sequence element, and (iii) making decisions on putative binding sites. The position weight matrix (PWM), (sometimes called position frequency matrix (PFM)) is the most commonly used model to represent the DNA binding preferences of TFs Stormo (2000). PWM is a 4 × *K* matrix where *K* is the expected length of the TF binding sites and each row of the matrix encodes one DNA nucleotide (A, C, G, T). Entries of the PWM represent the probability of nucleotides to appear at each position and the higher the value means the more conserved the DNA base at this position. Indeed, PWM assumes that the binding site nucleotides contribute independently to the binding specificity of TF. To relax this assumption, many researchers investigated the dependency amongst TFBS positions and its role in the DNA binding affinity of TF proteins Zhao *et al.* (2012); Quader and Huang (2012); ?. As a result, several alternatives for the classic PWM were proposed, e.g., di-nucleotide weight matrix Siddharthan (2010) and tree-based PWM Bi *et al.* (2011). However, recent work showed that the simple PWM well models the DNA binding specificities for most of the transcription factor proteins Zhao *et al.* (2012).

Computational methods usually scan the input DNA sequences using the PWM model of the investigated TF with a shifting window of width *K*. The K-mers produced by the shifting window are then characterized by some numerical values (features) that describe the binding affinity of the TF against each of these K-mers. Traditional computational approaches calculate scores for each K-mer using the information content of PWM and the probabilities of the K-mer bases at each position in the PWM, e.g., TESS Schug and Overton (1998), MatInspector Quandt *et al.* (1995), MATCH Kel *et al.* (2003). These methods solely depend on the K-mer DNA sequence to score the putative binding sites and do not use any prior knowledge on the putative TFBSs. Therefore, these approaches have limited performance since TFBSs are short and degenerate and occur spuriously in genomes. The TFBS bases by themselves cannot provide sufficient information about protein-DNA binding. To overcome this hurdle in the prediction of TFBSs, many researchers used the phylogenetic footprinting information from multiple sequence alignments of different species, e.g., MONKEY Moses *et al.* (2004) and Contra Hooghe *et al.* (2008). Other approaches used hidden Markov models (HMM) learnt on multiple genomes to model the phylogenetic footprinting, e.g., MAPPER Marinescu *et al.* (2005) and CONTRASIF Tokovenko *et al.* (2009). The motivation behind this is that functional DNA elements are evolutionary conserved across orthologous genome regions. Most of these approaches used PWMs from the TRANSFAC database Matys *et al.* (2006) and JASPAR Sandelin *et al.* (2004). Moreover, a special database was created for the common conserved motifs in the plant and chordate genomes and then a tool called DoOpSearch was developed to find putative TFBSs Sebestyen *et al.* (2009). The most recent work that used phylogenetic information to locate TFBS is ConTra v2 Broos *et al.* (2011) which allows users to search for putative TFBSs in any genomic region (promoters, 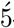-UTR, 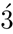-UTR or introns) for nine reference organisms. Contra v2 uses Multiz multiple sequence alignment Blanchette *et al.* (2004) to identify conserved regions and has an extensive library of curated PWMs. On the other hand, the 3D structure of the TF protein was used to improve finding the 3D structure of the DNA sequences of K-mers and their flanking regions Xu *et al.* (2013). More structural features can be calculated for K-mers to characterize functional TFBS from random background K-mers, e.g., PhysBinder computes some biophysical properties for the DNA bases to predict TFBSs Hooghe *et al.* (2012); Broos *et al.* (2013). Therefore, the precision of TFBS prediction is increased and the number of false positives is highly reduced. PhysBinder uses random forests Breiman (2001) trained on PWM and structural features to model the DNA binding affinity of TF proteins. Notwithstanding the usefulness of structural features, knowledge of them is still limited for most TF proteins.

The last step in TFBS prediction is to classify K-mers into putative binding sites or background non-functional sequences. All the state-of-the-art methods use a cut-off threshold to make the classification decision and their performance is very sensitive to this threshold. Most of the methods manually set different thresholds for each PWM so that they aim to reduce the number of false positives or increase the recall Cartharius *et al.*(2005); Kel *et al.* (2003). In this paper, we propose a threshold-free method, named FSCAN, that models the DNA binding affinity of TF proteins using fuzzy inference systems (FISs). FSCAN takes advantage of the available ChIP-chip data for TF proteins and combines them with phylogenetic information extracted from Multiz multiple sequence alignments in order to model TFBSs. Our system prediction flows into two stages:pre-processing and classification. In addition, a learning unit is developed to train a distribution-based filter using our adaptive ellipsoidal filter (AEF) algorithm and the adaptive neuro-fuzzy inference system (ANFIS) algorithm is used to extract fuzzy rules for the FIS. The AEF algorithm helps reduce the imbalance ratio between functional K-mers (TFBSs) and nonfunctional K-mers (background) in the training sequence-set. Furthermore, oversampling-based techniques are employed to reduce the imbalance ratio in the training dataset before it is introduced to the ANFIS learning algorithm. Our results clearly show that FSCAN is an efficient and effective solution for finding TFBSs in intergenic regions of probes bound by TFs in ChIP-chip experiments. Our method is tested on ChIP-chip data from the yeast genome and is compared with MatInspector and MATCH. FSCAN significantly outperforms these two methods and shows promising results for using fuzzy logic in identifying TFBSs.

The rest of the manuscript is structured into three sections as follows. Section 2 formulates the problem, reviews two popular methods for TFBS identification and reveals the challenges of the problem. Section 3 describes our proposed system FSCAN in detail. Section 4 analyzes the performance of FSCAN and investigates the robustness of our proposed algorithms. Section 5 concludes this work with observations on the proposed approach.

## 2 Related Work

### 2.1 Problem Statement

Given a set of intergenic sequences 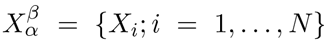 that are bound by the same transcription factor (TF) protein *β* under specific growth condition *α* in a ChIP-chip experiment and have different lengths. Our task is to find the locations of the DNA binding sites on which the TF protein may bind. It is assumed that the binding specificity of the TF protein is given as a position frequency matrix (PFM). The forward and reverse strands of each intergenic sequence are scanned so that each sequence strand of length *L* is partitioned into *L* − *K* + 1 K-mers (short sequences of lengths *K*; where *K* is the width of PFM). Finally, the K-mers that are most likely to be functional binding sites for the studied TF protein are reported.

In the next subsections, *X_ij_* represents a K-mer in sequence *X_i_* that starts at position *j* with length *K*. 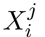 represents the nucleotide at position *j* in sequence *X_i_* where 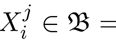 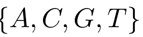. The binding specificity of TF proteins is represented by the position frequency matrix (PFM) *M_f_* or the position weight matrix (PWM) M_*w*_. The positions of DNA bases in the motif matrix (columns) are zero-based indexes and the width of the matrix is *K*. It is assumed that one of these two matrices is known for the TF proteins under study. It is worth mentioning that the entries of PFM are the bases’ frequencies at each position of the TF motif while PWM entries represent the log-odds probabilities of these frequencies.

### 2.2 MatInspector and MATCH

The idea behind the MatInspector Quandt *et al.* (1995) and MATCH Kel *et al.* (2003) programs is quite simple as they work in a similar way. Both programs scan the input sequence using the PFM model of a TF protein and assign two similarity scores for each K-mer:the core similarity score (CSS) and the matrix similarity score (MSS). CSS is calculated based on the most conserved and consecutive positions in the PFM model and is used to accelerate the scanning process while MSS is calculated based on all positions for K-mers that pass a cut-off threshold on CSS (see below). The only two differences between MatInspector and MATCH are the number of pre-computed PFM models in their databases and the way that they score the similarity of K-mers to PFMs. MatInspector had around 200 PFMs for different transcription factor proteins while MATCH used all the TF protein matrices (more than 500) that were available in the TRANSFAC database Matys *et al.*(2006) at the time of publication. CSS and MSS are calculated as follows.

#### 2.2.1 Matrix Similarity Score

This score expresses the relative similarity between a given K-mer *X_ij_* and a TF motif model represented by PFM *M_f_*. MSS is calculated based on the Shannon information content Schneider *et al.* (1986) of the individual positions of motif matrix. In MatInspector Quandt *et al.* (1995), it is defined as follows

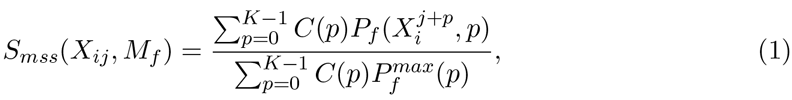

where 0 ≤ *S_mss_*(*X_ij_,M_f_*) ≤ 1, *P_f_* (*b*,*p*) is the probability of observing base *b* at position *p* from PFM, *C*(*p*) measures the conservation of bases at position *p* in PFM and 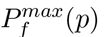 is the maximum nucleotide frequency at position *p*. Multiplying the base frequencies P*_f_* (*b*,*p*) by the values of *C*(*p*) emphasizes the fact that mismatches at lowly conserved positions are more easily tolerated than mismatches at highly conserved positions Quandt *et al.* (1995). Therefore, *C*(*p*) is calculated as follows

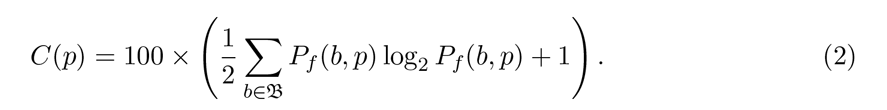

*C*(*p*) equals 100 when position *p* is very conserved (i.e., the same nucleotide appears all time) and equals 0 when position *p* is not conserved at all (i.e., all nucleotides appear equally). On the other hand, MATCH Kel *et al.* (2003) calculates MSS using a slightly different formula, that is

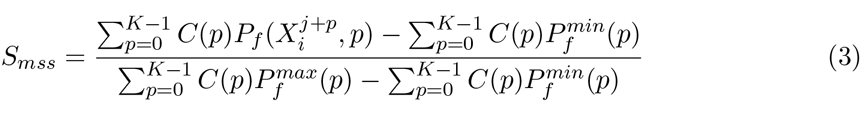

where the conservation vector *C*(*p*) is defined differently from Eq. 2, as follows

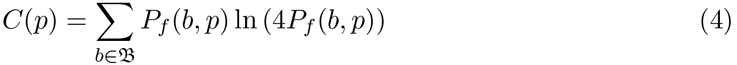

#### 2.2.2 Core Similarity Score

CSS is calculated only based on the most conserved core region within the PFM matrix Quandt *et al.* (1995); Kel *et al.* (2003). A core region is the 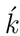 consecutive nucleotide positions with the highest sum of *C*(*p*) values where 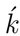 is usually set to 4 or 5. In MatInspector, CSS is defined as follows

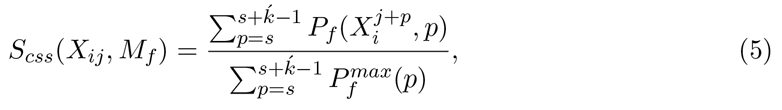

where 0 ≤ *S_cSS_*(*X_ij_,M_f_*) ≤ 1 and 0 ≤ *s* ≤ *K* – 1 is the start position of the core region. MATCH, however, calculates the CSS using Eq. 3 with only the core region positions in the motif matrix.

### 2.3 Challenges

After assigning the matrix similarity scores (MSSs) to K-mers, a decision needs to be made as to whether a K-mer is a putative binding site or non-functional background sequence. In other words, a cut-off threshold should be set to make this decision and this is one of the biggest challenges for the threshold-based TFBSs prediction systems. These systems are very sensitive to the cut-off threshold value and should be selected carefully. They only use similarity between a K-mer and a motif matrix to predict binding sites and no prior knowledge on the K-mer sequence is applied. To illustrate the threshold setting problem in MatInspector and MATCH, both programs are executed at different cut-off thresholds between 0.85 and 1.0 on 22 ChIP-chip datasets (dataset details are explained in Section 4.1). Table 1 shows the average performance indexes (see Section 4.2) of MatInspector over all datasets and at different cut-off thresholds *θ* It is obvious how the cut-off threshold influences the performance of the system. A slight change in the cut-off threshold results in a big difference in the number of predicted binding sites. To further understand the effect of the cut-off threshold on each dataset independently, Fig. 1 was plotted. Fig. 1 illustrates the average F1-Measure and Performance Coefficient (PC) for MatInspector on each dataset for cut-off thresholds in [0.85,1.0]. The standard deviation of these measures is also calculated and depicted for each dataset (vertical lines on Fig. 1). It can be easily seen how the setting of this threshold can dramatically change the system performance for some datasets (standard deviation is larger than 0.1). Fig. 1 also shows the performance of MatInspector at the best performing cut-off threshold for each dataset. F1 and PC at the best performing threshold represent the cut-off threshold that minimizes both the number of false negatives and the number of false positives. Similar observations are seen in the MATCH method. This clearly shows the limitations of the MatInspector and MATCH approaches. Some datasets could not perform more than 0.4 for F1 (less than 0.3 for PC) due to the heavy overlapping between functional K-mers and background K-mers. In other words, the same DNA sequence appears functional in some locations and non-functional in others.

**Figure 1:**
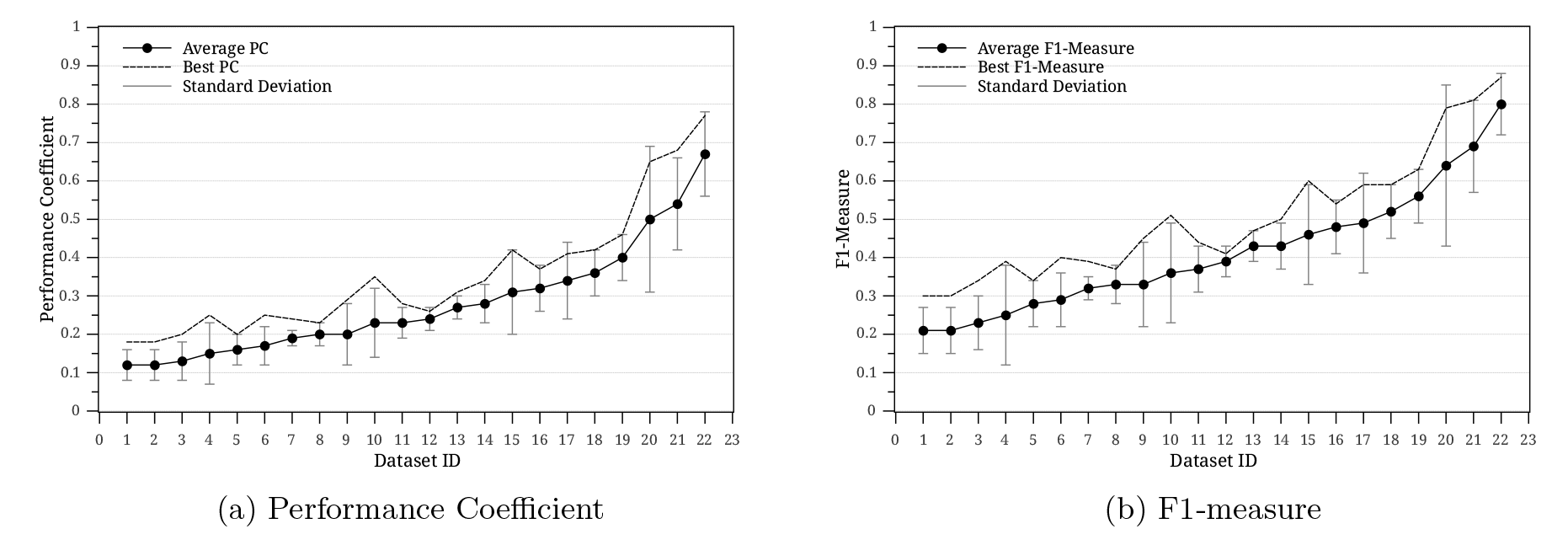
The performance of MatInspector at different cut-off threshold values in [0.85, 1].

**Table 1:**
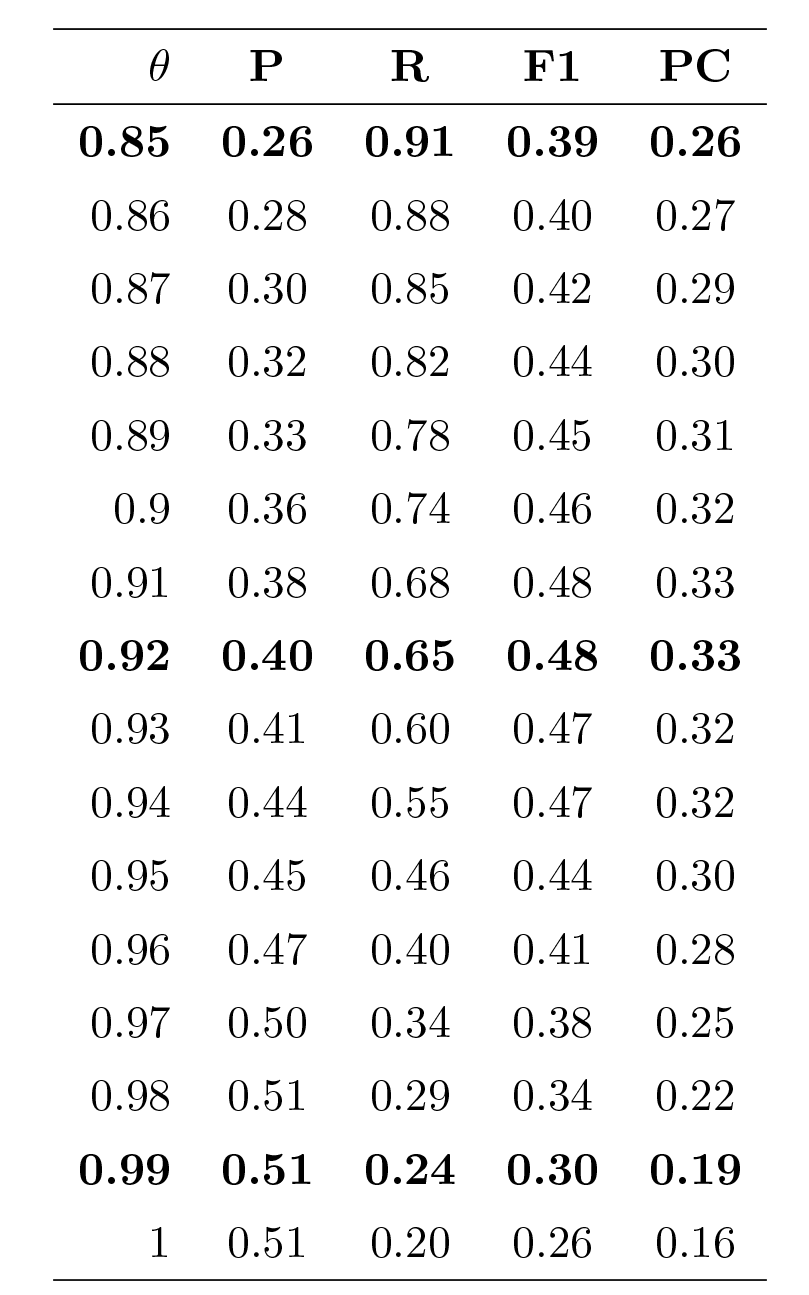
The average performance of MatInspector on all datasets and for different cut-off thresholds.

| $\theta$ | P | R | F1 | PC |
| --- | --- | --- | --- | --- |
| **0.85** | **0.26** | **0.91** | **0.39** | **0.26** |
| 0.86 | 0.28 | 0.88 | 0.40 | 0.27 |
| 0.87 | 0.30 | 0.85 | 0.42 | 0.29 |
| 0.88 | 0.32 | 0.82 | 0.44 | 0.30 |
| 0.89 | 0.33 | 0.78 | 0.45 | 0.31 |
| 0.9 | 0.36 | 0.74 | 0.46 | 0.32 |
| 0.91 | 0.38 | 0.68 | 0.48 | 0.33 |
| **0.92** | **0.40** | **0.65** | **0.48** | **0.33** |
| 0.93 | 0.41 | 0.60 | 0.47 | 0.32 |
| 0.94 | 0.44 | 0.55 | 0.47 | 0.32 |
| 0.95 | 0.45 | 0.46 | 0.44 | 0.30 |
| 0.96 | 0.47 | 0.40 | 0.41 | 0.28 |
| 0.97 | 0.50 | 0.34 | 0.38 | 0.25 |
| 0.98 | 0.51 | 0.29 | 0.34 | 0.22 |
| **0.99** | **0.51** | **0.24** | **0.30** | **0.19** |
| 1 | 0.51 | 0.20 | 0.26 | 0.16 |

## 3 Fuzzy Prediction of TFBS Locations (FSCAN)

It is obvious from our previous argument that identifying transcription factor binding sites using traditional threshold-based systems relies more on handcrafting than science. Therefore, we propose a threshold-free system that integrates prior phylogenetic knowledge with sequence information in order to locate the binding sites of a particular TF protein in intergenic sequence-sets. Instead of using crisp logic as in traditional methods, our system, named FSCAN, uses fuzzy inference techniques to decide whether a K-mer is a putative binding site or non-functional sequence. This gives our system a good ability to detect weak binding sites that are similar to non-functional K-mers. In order to generate the fuzzy rules that are used in the inference system, an adaptive neuro-fuzzy system (ANFIS) neural network was used along with two training and validation sequence-sets. FSCAN is composed of three main components that interact together to produce accurate predictions of TFBSs. These three components are:pre-processing, learning and classification units. Each component performs a set of functionalities as detailed next. Fig. 2 illustrates the FSCAN components and functionalities.

**Figure 2:**
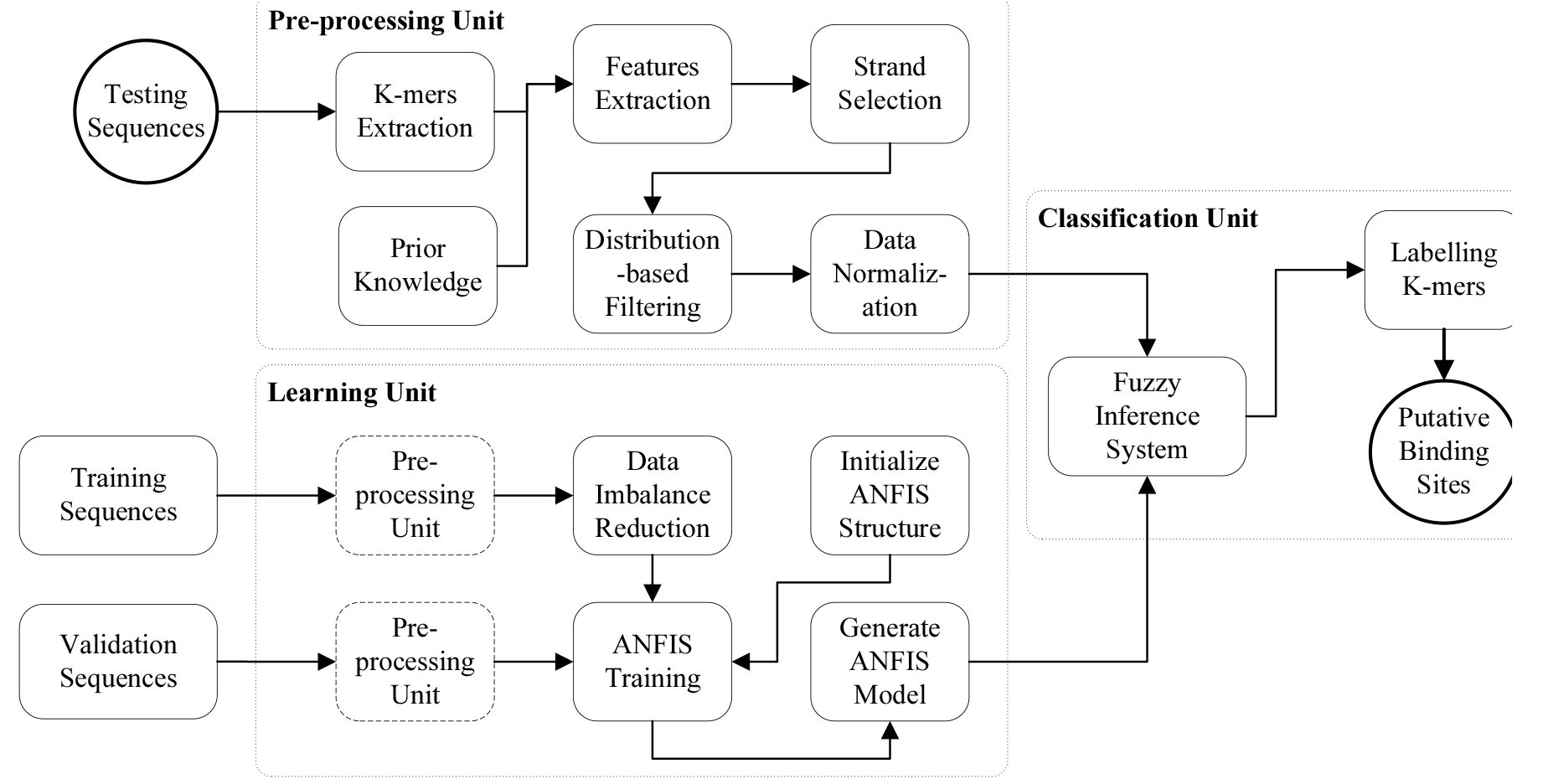
FSCAN system components and functionalities.

### 3.1 Pre-processing Unit

In this unit, the intergenic DNA sequences are initialized to be fed into the system for testing or learning. K-mers are extracted from these sequences and the ones that are less likely to be putative TFBSs are removed in this step and only the locations that could contain binding sites remain by the end of the pre-processing stage. Moreover, features are extracted to model TFBSs. Four main functionalities are included in this unit, as follows.

#### 3.1.1 K-mers Extraction

This function partitions DNA sequences into K-mers (short sequences of length *K* where *K* is the TF motif width). A window of size *K* is used to scan DNA sequences and generate K-mers by shifting the window one nucleotide position each time. As a result, a sequence of length *L* is partitioned into *L* – *K* + 1 K-mers. For example, the following portion of an intergenic sequence *ATGTACTAGAATGTGATGGAGTGGGGGTT* in *ABF* 1_*YPD* sequence-set produces these 13-mers:*ATGTACTAGAATG, TGTACTAGAATGT*, *GTACTAGAATGTG,…,TGGAGTGGGGGTT*. This procedure is applied similarly on the two DNA strands of intergenic regions.

#### 3.1.2 Features Extraction

In order to discriminate true binding sites from other background sequences, three features are extracted for each K-mer. The first two features are calculated using MISCORE Wang and Tapan (2012) and are based on the motif model (PFM). They are denoted by the mean and width of the TF binding affinity to a given DNA sequence. On the other hand, the third feature solely depends on the location of the K-mer in the genome and its conservation across other related species. We call it simple phylogenetic footprinting. It is obvious that calculating these three features requires prior knowledge of the PFM of the TF protein and the multiple sequence alignment of orthologous sequences. However, we introduce the similarity metric MISCORE first. Mismatch score (MISCORE) Wang and Tapan (2012) was published as a scoring function in order to distinguish functional TF motifs from random background motifs in the de novo motif discovery applications. Furthermore, it can be used to measure the similarity between a given K-mer and a PFM model. It takes into account the similarity between the K-mer and the background sequences (described with a PFM) in addition to the similarity between the K-mer and the TF motif matrix. This metric relaxes any assumption on the position dependency and uses a simple approach to represent background reference model *M_ref_*. MISCORE between a K-mer *X_ij_* and a PFM *M_f_* is calculated by

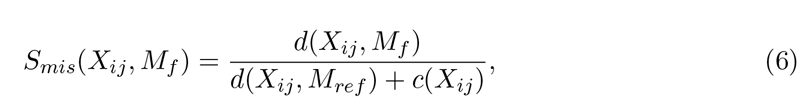

where *S_mis_*(*X_ij_,M_f_*) ≥ 0 (the smaller the value, the more probable it is to be a binding site), *M_ref_* is the background reference model as a PFM calculated using all possible K-mers in the intergenic sequence-set, *d*(*X_ij_,M_f_*) is the generalized Hamming distance between a K-mer encoding matrix and a PFM and is expressed as

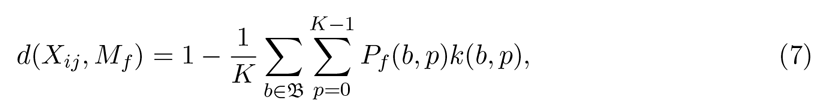

where *k* is a 4 × *K* binary matrix encoding the K-mer *X_ij_* defined as follows

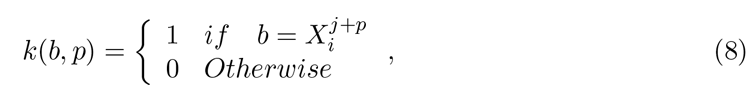

and *c*(*X_ij_*) in Eq. (6) represents the compositional complexity of a K-mer and is quantified using the distribution of nucleotides in the K-mer, as follows

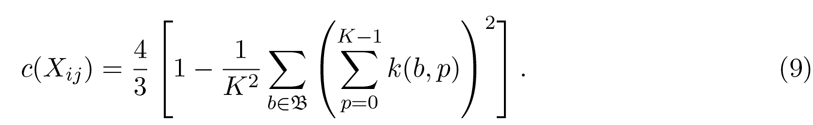

However, ChIP-chip experiments can produce two sequence-sets based on the binding probability of TF to a probe (*p-value*). A bound sequence-set, denoted by *X*, contains the DNA sequences that are most likely to be bound by the TF protein (usually having a very low *p-value*) and the unbound sequence-set, denoted by *Y*, contains the sequences that are least likely or never bound by the TF protein during the experiment (usually having a very high *p-value*). Therefore, a modified MISCORE function, named discriminative conserved MISCORE (DC-MISCORE), is defined. It differs from the original MISCORE in Eq. (6) in two aspects. First, it is more discriminative, that is K-mers that appear in the bound sequence-set less frequently than they appear in the unbound sequence-set have higher scores. Second, it takes into account the conservation of each position in the PFM, that is highly conserved positions contribute significantly in the K-mer mismatch score while lowly conserved positions have less impact on the final score. As a result, the DC-MISCORE is defined using this formula

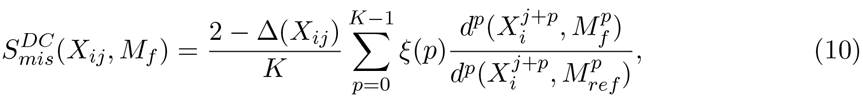

where 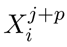, 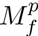, and 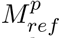 are the *p*th positional columns in *X_ij_*, *M_f_*, and *M_ref_*, respectively, *d*()^*p*^ is a special generalized Hamming distance that measures the dissimilarity between a motif PFM and a K-mer sequence at a specific position, as follows

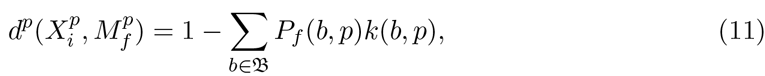

ξ(*p*) is the degree of conservation of position *p* in the *M_f_* matrix and is given by the

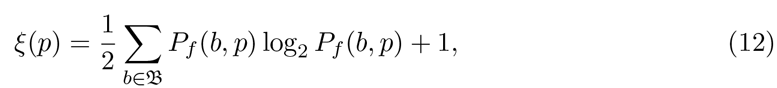

and Δ(*X_ij_*) represents the over-representation of K-mer *X_ij_* in the bound sequence-set *X*

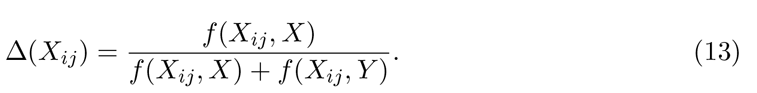

where *f*(*z*, *Z*) is the frequency of K-mer *z* in the sequence-set *Z*.

In the next section, an explanation of how to calculate the three features that ease the modelling of TFBSs is given.

**Mean and Width of Binding Affinity** The strength of DNA-protein binding strongly depends on which amino acids in the protein contact which DNA nucleotide in the DNA sequence Luscombe and Thornton (2002). However, some amino acids in the DNA-binding domain of TF proteins strongly bind to the binding site bases while others weakly bind to their corresponding DNA bases Zhao *et al.* (2012). This motivated us to define the similarity between a K-mer and a motif PFM so that not all positions in the PFM and the K-mer sequence are included in calculating the similarity score. Different from MatInspector Quandt *et al.* (1995) and MATCH Kel *et al.* (2003), which used the concept of core region, we propose a novel methodology to estimate the binding affinity between a TF protein and a putative binding site. *R* positions are selected randomly from the PFM of the TF protein with their corresponding bases in the K-mer sequence. Then, the similarity between the K-mer and the PFM is calculated based on the randomly selected positions *R_p_* only, using a modified formula of Eq. (10) as follows

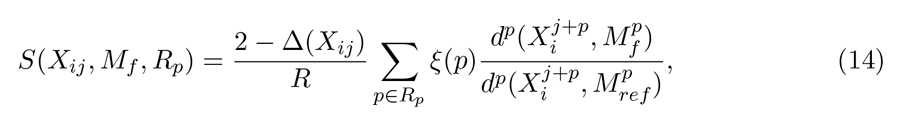

where Δ(*X_ij_*) is the K-mer over-representation ratio as given in Eq. (13), ξ(*p*) is the PFM conservation vector as defined in Eq. (12), and *d*^*p*^() is the Hamming distance as defined in Eq. (11). This random sampling of positions is repeated *T* times and S(*X_ij_, M_f_,R_p_*) is calculated at each trial. A *DNA binding affinity signal* (DNA-BAS) is generated for each K-mer with respect to the studied TF protein. As a result, K-mers that could be putative binding sites will have quite low amplitude for their DNA-BASs while non-functional K-mers will have very high amplitude, as depicted in Fig. 3a.

**Figure 3:**
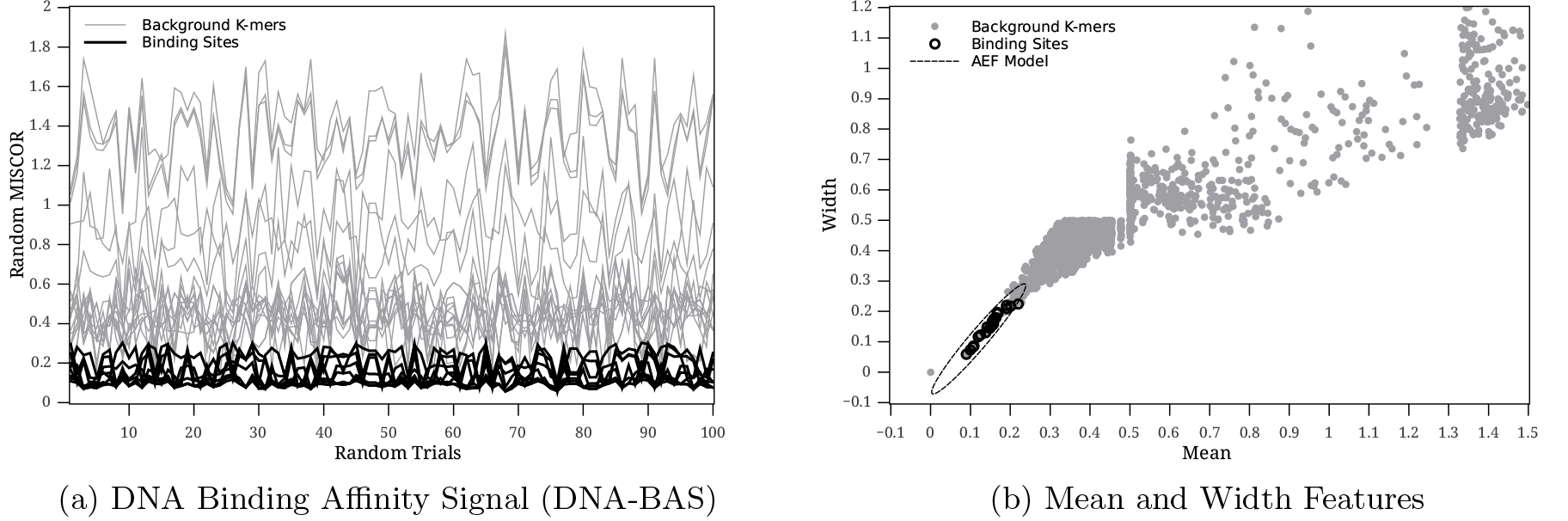
The DNA binding affinity of UME6 YPD protein for non-functional K-mers and true binding sites.

Two features can now be extracted for each K-mer from the DNA-BAS, namely, the *mean* of binding affinity which is calculated using the following formula

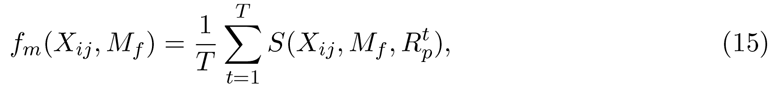

where 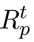 is the set of randomly selected positions at trial *t*, and the *width* of the binding affinity, i.e., the peak-to-peak amplitude of the binding affinity signal which is defined as

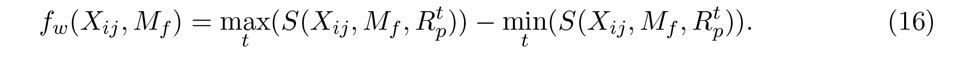

The number of selected positions *R* is usually set to 60% of the motif width, that is *R* = [0.6 × *K*] and the number of random trials *T* is usually set to 100. Fig. 3b illustrates the mean and width features for K-mers extracted from the bound sequence-set of *UME*6_YPD ChIP-chip data.

**Simple Phylogenetic Footprinting** This feature determines at which level a potential binding site is conserved across different species. The rationale behind this feature is that genomic regions that contain TFBSs have very strong phylogenetic relationships Blanchette and Tompa (2002). For this purpose, the conservation scores for the S. Cerevisiae are obtained from the UCSC genome browser. These scores are produced for each base in the genome using phastCons program. phastCons Siepel *et al.* (2005) gives the posterior probability of a nucleotide position being generated by the conserved state of a two-state phylogenetic hidden Markov model (phylo-HMM). Phylo-HMM takes into account how nucleotides alternate at each position and how this process changes from one position to the next. The phylo-HMM model is fitted on a 7-way MULTIZ Blanchette *et al.* (2004) multiple sequence alignment by maximum likelihood using the expectation-maximization (EM) algorithm. This multiple sequence alignment is formed from the alignment of 6 yeast genomes with the S. Cerevisiae genome:*S. Cerevisiae*, *S. Paradoxus*, *S. Mikatae*, *S. Kudriavzevii, S. Bayanus, S. Castelli* and *S. Kluyveri*. To convert these conservation scores into a discriminative feature that characterizes TFBSs, a simple phylogenetic footprinting score is calculated for each K-mer *X_ij_*, as follows

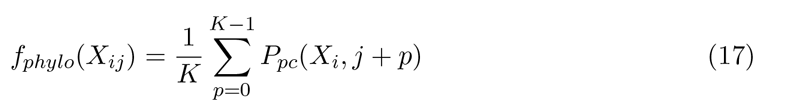

where *P_pc_*(*X_i_; p*) is the phastCons score at position *p* in sequence *X_i_*. This feature plays a key role in discriminating functional binding sites from non-functional K-mers that have the same DNA nucleotides sequence but occur in other genomic regions.

Likewise, more features can be used to better model true binding sites. However, it is out of the scope of this paper to investigate such features. By the end of this step, each K-mer in the sequence-set is encoded by a 3-dimensional features vector (f_m_; f_w_; f_phylo_) ∈ ℝ^3^ and a dataset of vectors is constructed.

#### 3.1.3 K-mer Filtering

Since we are dealing with very large-scale sequence-sets, a two-stage filtering procedure is designed to reduce the number of non-functional background K-mers in the sequence-sets. In the first stage, one of the two strands (forward and reverse) for each K-mer is selected to be in the training/testing sequence-sets. We filter out the K-mers that have larger mean values *f_m_* than their counter K-mers on the other strand. For example, if the mean value of 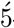 — *ACGCTG* — 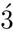 is 0.35 and the mean value of its reverse complement on the other strand 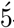 — *CAGCGT —* 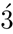 is 0.2, then the forward K-mer is stripped out. If the two K-mers have equal means of binding affinity *f_m_* (i.e., palindromic sequence), then the K-mer on the reverse strand is filtered out.

The second stage of filtering removes all K-mers that fall outside an ellipsoidal region that surrounds the true binding sites (see Fig. 3b). The mean *f_m_* and width *^f^w* of the binding affinity features are used to design the filter. In order to define a boundary ellipse for the true binding sites in the 2-dimensional feature space *(f_m_,f_w_*) ∈ ℝ^2^, we develop our own algorithm based on principle components analysis (PCA) Pearson (1901). First, 2D feature vectors are constructed from the 3D feature vectors dataset (usually training and validation datasets) and using the K-mers of the *true binding sites* only. Then, they are organized in a matrix of *N* × 2 elements, denoted by *F*. Second, the means of these two features over all vectors in *F* are calculated, as follows

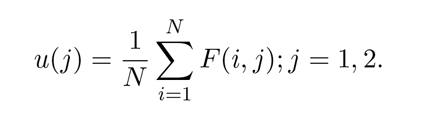

Third, all data points are centered around the mean *u* as follows

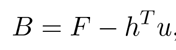

where *h* is a 1 × *N* vector whose entries are all equal to 1. Fourth, a 2 × 2 covariance matrix *C* is found for the matrix *B* using this formula

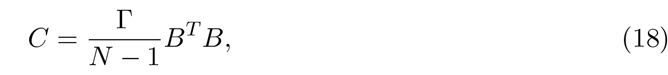

where Γ ≥ 1 is a scaling coefficient to control the filter sensitivity. Then, the eigenvectors of *C* that transform *C* into a diagonal matrix *D* of the eigenvalues of *C* are computed and stored in a 2 × 2 matrix *V*, as follows

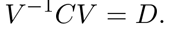

Eventually, the basis of the eigenvectors space is scaled by the square root of the corresponding eigenvalues as follows

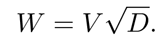

Now, the ellipsoidal filtering procedure is applied on all K-mers to filter out irrelevant background K-mers in three steps

- center the features vector (*f_m_, f_w_*) around the mean *u*.
- project the centered vector on the new basis 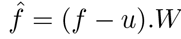.*W*
- test the ellipsoidal filtering equation

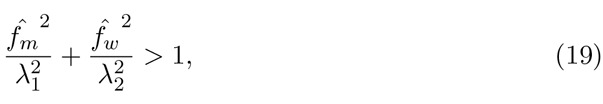

where λ_1_ and λ_2_ are the eigenvalues of the covariance matrix *C*.
- K-mers whose mean and width features satisfy Eq. (19) are removed.

**Algorithm 1.**
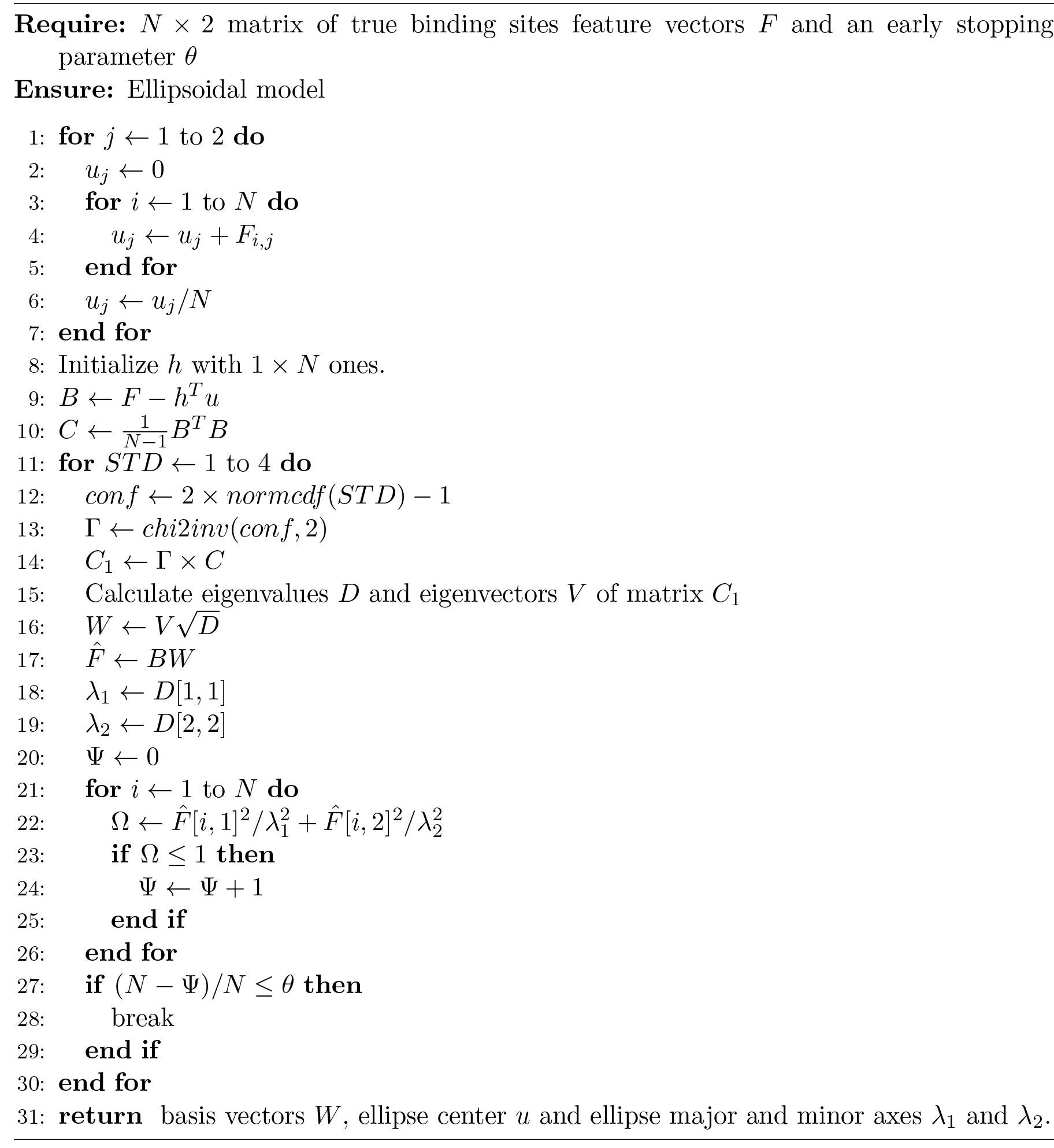
Adaptive ellipsoidal filter (AEF) algorithm.

During the filter design, the sensitivity parameter Γ. in Eq. (18) should be set carefully so that the filter does not remove many true binding sites. Multiplying the covariance matrix *C* with a scaling factor is equivalent to multiplying the eigenvalues of the matrix by this factor. Moreover, these eigenvalues represent the lengths of the major and minor axes of the ellipsoidal region. In other words, Γ determines how much of the K-mers are within a certain standard deviation away from the mean point. Inverse-chi-square distribution is adopted in order to estimate Γ that covers some percentile of the data with two degrees of freedom (because there are two features). Here it is assumed that the two random variables corresponding to the features (*f_m_, f_w_*) are normally distributed and hence Eq. (19) follows a chi-square distribution. Therefore, 68% of confidence interval corresponds to 1 standard deviation away from the mean, 95% corresponds to 2 standard deviations away from the mean, and so on. Since each dataset has different K-mers distribution in the feature space, an adaptive algorithm is proposed to estimate the value of Γ for each sequence-set separately.

Algorithm 1 shows how the adaptive ellipsoidal filter (AEF) is learnt. In Algorithm 1, *normcdf (std)* is the cumulative distribution function (CDF) of the normal distribution and returns the area under the normal density curve from −∞ to *std* Therefore, 2 × *normcdf (std) –* 1 returns the confidence interval that corresponds to std standard deviation away from the mean. *chi2inv*(*conf*, 2) computes the inverse of the chi-square CDF with two degrees of freedom and returns the value which exceeds *conf* * 100% of the samples from a chi-square distribution with 2 degrees of freedom. Eventually, the error acceptance level *θ* controls the number of background K-mers that will be filtered in and helps in the early stopping of the learning algorithm. 0.02 is the default value for this parameter.

Table 2 shows the impact of the ellipsoidal filtering on the investigated sequence-sets. It greatly reduces the number of non-functional K-mers. BG/BS measures the ratio of the number of background K-mers to the number of known binding sites in the sequence-set and is called the imbalance ratio. The out BS% column shows the percentage of true binding sites that are filtered out because of the K-mer filtering procedure. It can be easily seen that this pre-processing step greatly reduces the imbalance ratio between functional and non-functional K-mers. The total imbalance ratio is reduced from 1940 to only 20 and the total number of extracted K-mers drops from 4 × 10^6^ to 4 × 10^4^ with almost 99% overall accuracy.

**Table 2:** The size of datasets before and after the filtering functionality.

| Dataset | Before Filtering |  | After Filtering |  |  |
| --- | --- | --- | --- | --- | --- |
|  | BG/BS | Size | BG/BS | Size | Out BS% |
| ABF1_YPD | 1 155 | 174 602 | 1 | 228 | 0.66 |
| CBF1_SM | 2 032 | 239 840 | 2 | 396 | 1.69 |
| CIN5_YPD | 4 741 | 275 034 | 34 | 1 932 | 3.45 |
| DIG1_BUT14 | 1 104 | 88 396 | 35 | 2 844 | 1.25 |
| FHL1_YPD | 3 240 | 200 930 | 13 | 812 | 3.23 |
| FKH1_YPD | 2 424 | 140 670 | 9 | 565 | 1.72 |
| GCN4_SM | 1 988 | 198 894 | 5 | 636 | 1 |
| HAP1_YPD | 3 200 | 188 844 | 44 | 2 667 | 0 |
| MBP1_H2O2Hi | 1 139 | 115 148 | 2 | 285 | 2.97 |
| NDD1_YPD | 2 568 | 143 842 | 64 | 3 591 | 1.79 |
| NRG1_H2O2Hi | 4 125 | 251 674 | 7 | 517 | 0 |
| PHD1_BUT90 | 1 489 | 235 394 | 11 | 1 822 | 0 |
| RAP1_YPD | 2 589 | 170 936 | 13 | 901 | 3.03 |
| REB1_H2O2Lo | 1 346 | 184 552 | 1 | 208 | 2.19 |
| SKN7_H2O2Lo | 2 381 | 266 758 | 52 | 5 803 | 1.79 |
| SOK2_BUT14 | 2 119 | 171 722 | 44 | 3 650 | 0 |
| STE12_Alpha | 2 385 | 248 180 | 23 | 2 522 | 0 |
| SUT1_YPD | 1 328 | 179 398 | 41 | 5 635 | 0 |
| SWI4_YPD | 1 604 | 227 874 | 8 | 1 331 | 0 |
| SWI5_YPD | 1 734 | 159 610 | 27 | 2 561 | 0 |
| SWI6_YPD | 1 185 | 190 948 | 6 | 1 069 | 0 |
| UME6_YPD | 2 046 | 131 034 | 1 | 134 | 1.56 |
| **Summary** | 1 940 | 4 184 280 | 20 | 40 109 | 1.02 |

#### 3.1.4 Data Normalization

Because of the filtering step, the scale of K-mer features changes according to the filtering conditions. Therefore, all features of the remaining K-mers are normalized to have the same scale [0,1] using this formula

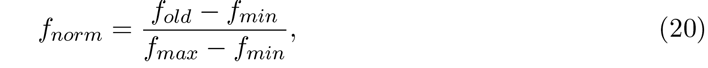

where *f_old_* and *f_norm_* are the feature value before and after the normalization, respectively, and *f_min_* and *f_max_* are the minimum and maximum feature values, respectively.

### 3.2 Learning Unit

The same pre-processing criteria and functionalities are applied on the training and validation sequence-sets in order to generate training and validation datasets (see Fig. 2). In addition, a K-mer is labelled with +1 if its location *exactly* matches a location of known binding site and with –1 otherwise. As a result, two datasets of labelled data pairs (*f,c*) ∈ ℝ^3^ × {+1, –1{ are created. The training and validation datasets are then used by the ANFIS learning algorithm to train the ANFIS neural network and thus generate fuzzy sets and fuzzy rules to be used in the fuzzy inference system (FIS). However, two steps are needed to generate an efficient FIS for identifying TFBSs.

#### 3.2.1 Data Imbalance Reduction

(It is obvious from (Table 2 that the imbalance ratio BG/BS) is still large for most of the datasets although a filtering procedure is applied. This leads learning algorithms to be biased towards the majority class data (the background K-mers in FSCAN) and unable to predict the minority class examples (true binding sites). Indeed, the imbalanced data problem causes most machine learning algorithms to fail to learn the underlying data distribution He and Garcia (2009). We use two techniques in order to reduce the imbalance ratio between functional and non-functional K-mers and alleviate this hurdle. The first one is based on random oversampling for the minority class He and Garcia (2009) and the second one is the well-known SMOTE algorithm Chawla *et al.* (2002).

Our minority oversampling algorithm, named random oversampling (RANDOVER), is simple and efficient. First, the ratio between the number of non-functional K-mers and the number of functional K-mers (BG/BS) is calculated. The oversampling technique isapplied on the training dataset iif the ratio is greater than 1. Then, the number of minority examples to be added to the imbalanced dataset is calculated. Next, Σ feature vectors are randomly sampled from the training dataset. Eventually, the replicated feature vectors are randomly perturbed with small noise in [−*δ*, +*δ*] and added to the original training dataset.The feature vectors in the new training dataset are then randomly shuffled before it is introduced to the learning algorithm of ANFIS. After applying RANDOVER on the training dataset, the imbalance ratio is reduced to less than 4 in most of the cases. Algorithm 2 explains the RANDOVER procedure. In our FSCAN system, *β* and *δ* are set to 0.25 and 0. 02, respectively.

**Algorithm 2.**
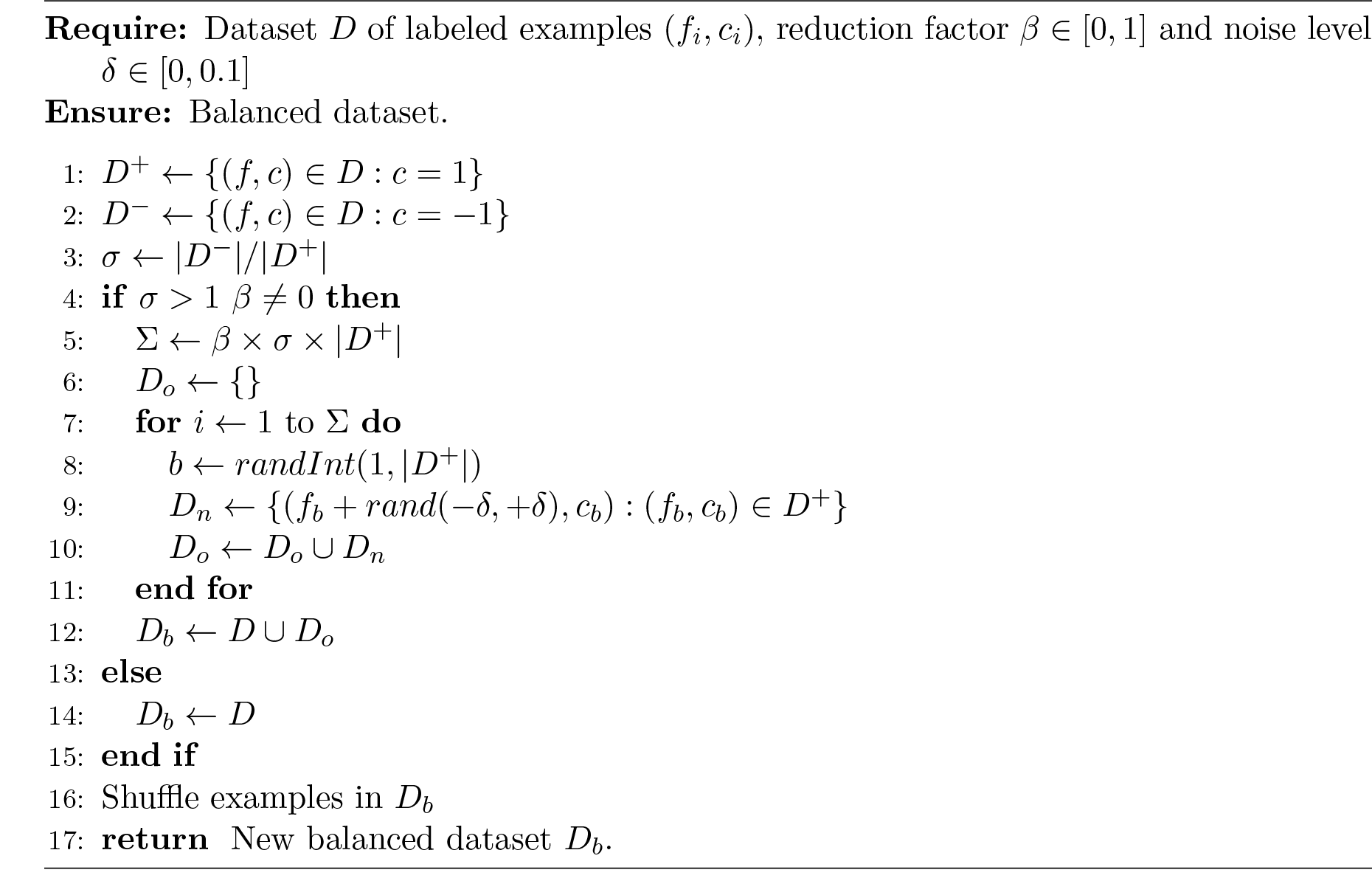
RANDOVER algorithm.

On the other hand, we apply the synthetic minority oversampling technique (SMOTE) on training sequence-sets to compare with our RANDOVER algorithm. SMOTE reduces the imbalance between minority examples (binding sites) and majority examples (background K-mers) by creating new artificial examples. The synthesized examples are created using each example and its nearest neighbor examples in the minority dataset, as follows

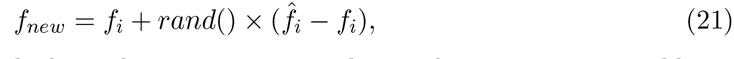

where *f_i_* and 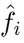 are an example from the minority set and one of its K-nearest neighbors, respectively. The new synthetic example lies somewhere on the line between *f_i_* and 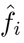 depending on the random coefficient *rand*(). For our FSCAN system, 4 nearest neighbors are selected for each minority example and then 3 new examples are synthesized for each.

#### 3.2.2 Generation of Fuzzy Rules

Fuzzy rules are the main component in any fuzzy inference system and linguistic variables are the essence of these rules. Each linguistic variable is expressed by several fuzzy sets and membership functions. The parameters of the membership functions and the fuzzy rules can be defined manually when expert knowledge is available. However, this is not the case in most applications, including our FSCAN system. Several techniques were proposed to extract fuzzy rules from data Jang (1993); Yongfu Wang *et al.* (2011). In this paper, the adaptive neuro-fuzzy inference system (ANFIS) is used Jang (1993) to define membership functions and generate fuzzy rules based on the training and validation features vectors/labels pairs. ANFIS is a special neural network composed of 6 layers of neurons and is not fully connected (details on the network architecture can be found in Negnevitsky (2001)). ANFIS can be used with the Takagi-Sugeno inference model only and has a single output variable that can be a constant (zero-order model) or linear (first-order model). ANFIS works in two stages. It first initializes a fuzzy inference system (FIS) with linguistic variables, fuzzy sets, membership functions and fuzzy rules. Then, it applies a combination of the least-squares method and the back-propagation gradient descent method to train FIS membership function parameters to emulate a given training dataset. A validation dataset can be also used to avoid model overfitting as it is the case in this paper.

As for our FSCAN system, four linguistic variables are defined for the system:three for the input (one for each K-mer feature) and one for the output (the K-mer class label). The universe of discourse for all input variables is [0,1] and three fuzzy sets (small, medium, large) are defined for each one. The generalized bell-shaped membership function is used to assign a grade of membership for each input to each fuzzy set. It is defined as follows

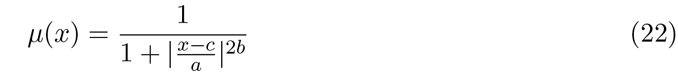

where *a,b,c*, and *d* are the membership function parameters that are learnt during the ANFIS training. Since there are only three input variables for FSCAN and each with three membership functions, the initial Sugeno FIS is generated from the training data using grid partitioning. Grid partitioning initializes the parameters of the membership functions so that the domain of each input variable is divided equally with sufficient overlapping. Then, it generates an initial rule base by enumerating all possible combinations of membership functions of all input variables, i.e., 27 rules are generated for our FSCAN system. The zero-order Sugeno fuzzy model is used in FSCAN, i.e., the consequences of rules are represented by singletons. After this, the training algorithm of ANFIS is run for 50 epochs. After training is completed, a rule base is obtained to be used later in the classification unit.

### 3.3 Classification Unit

In this unit, the Sugeno inference model Sugeno (1985) is applied on the learnt FIS in order to predict the locations of binding sites. K-mer features are introduced into the FIS which fuzzifies their crisp values and assigns degrees of membership to each feature value according to its corresponding fuzzy sets. For example, if a K-mer is described by the feature vector (0.1, 0.2, 0.9), then the degrees of memberships are calculated using Eq. (22) for each input feature and for all fuzzy sets (small, medium, large). Next, the fuzzified values are used to trigger the fuzzy rules and calculate their output values using the fuzzy operations. Subsequently, the outputs of all rules are aggregated and generate one single set of singletons. Finally, the weighted average of these singletons is calculated to defuzzify the output fuzzy value and convert it into a meaningful output for FSCAN. More details on the fuzzy inference can be found in Negnevitsky (2001).

The fuzzy inference system always produces real numbers. However, each K-mer must be labelled with a class label at the output of FSCAN. Consequently, a K-mer is classified as a true binding site if the FIS output is greater than 0 and as a background sequence otherwise. Moreover, all K-mers that are filtered out in the pre-processing unit are labelled as background sequences.

## 4 Performance Evaluation

The 10-fold cross-validation procedure is conducted to evaluate the performance of FSCAN on each ChIP-chip sequence-set. Each sequence-set is divided into 10 subsets so that each subset contains an equal number of background K-mers and true binding sites. The 10 subsets initiate 10 runs for each sequence-set in which each subset is tested once and used more than once for training or validation. For each run, two subsets are selected for testing and validation and the remaining subsets are combined together to make the training sequence-set. Then, the AEF algorithm (Algorithm 1) is invoked on the training and validation of K-mers to learn the ellipsoidal filter parameters. Afterwards, the filter is applied on the training, validation and testing of K-mers to label background K-mers. The remaining K-mers in the training and validation subsets are used to prompt the learning unit and generate the fuzzy inference system (FIS). Eventually, the remaining K-mers in the testing subset are classified using the generated FIS and are thus labelled. A 2 × 2 confusion matrix is created based on the labelled testing K-mers. The 10 confusion matrices produced from the 10 runs of cross-validation are summed together in order to calculate the evaluation metrics of FSCAN on one final confusion matrix.

On the other hand, the performance of MatInspector Quandt *et al.* (1995) and MATCH Kel *et al.* (2003) is assessed differently. Both programs are run using many values for the cut-off thresholds and for each sequence-set, i.e., all possible values in [0.85,1.0] are tested with an incremental step of 0.01. For each value, a confusion matrix with its evaluation metrics are calculated. Finally, the best performance measures of all based on F1-measure and Performance Coefficient (PC) are reported.

### 4.1 Datasets

To demonstrate the importance of our proposed algorithm, we collected ChIP-chip data published by Harbison *et al.* (2004) for 203 verified transcription factors from the Saccharomyces Cerevisiae budding yeast genome. The authors profiled the binding locations of these TFs over around 6000 intergenic regions (IGRs) that cover the whole yeast genome. The binding of each TF was investigated under at least one of the following 14 growth conditions:YPD (rich medium), Acid (acidic medium), Alpha (alpha factor pheromone treatment), BUT14 (butanol treatment for 14 hours), BUT90 (butanol treatment for 90 minutes), GAL (galactose medium), H2O2Hi (highly hyperoxic), H2O2Lo (mildly hyperoxic), HEAT (elevated temperature), Pi- (phosphate deprived medium), RAFF (raffinose medium), RAPA (nutrient deprived), SM (amino acid starvation), or THI- (vitamin deprived). As a result, 350 ChIP-chip experiments were run and each experiment produced one sequence-set of the form 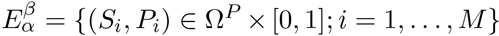, where *P_i_* is binding probability (*p—value*) of transcription factor *β* to probe sequence *S_i_* under growth condition *α* and *M* is the total number of probe sequences. Later, these sequence-sets were reanalyzed and an improved regulatory map was generated MacIsaac *et al.* (2006). In the new map, the motif matrices of 124 TFs were determined with their binding sites under the most stringent binding value *p — value* < 0.001 and under high confidence of conservation criteria (conserved in at least two other yeast species). The compiled TFBSs in the later study have been adopted by the Saccharomyces Genome Database (SGD) Cherry *et al.* (2012). Of these 124 transcription factors, 88 had well-known motifs in the literature at the time the work was published.

Moreover, the S. Cerevisiae genome, version *R*64.1.1 (published in February 2011), is downloaded from the Saccharomyces Genome Database (SGD) Cherry *et al.* (2012). Since the ChIP-chip experiments in Harbison *et al.* (2004) were applied on an old release of the yeast genome, the start and end positions of the probes and intergenic regions are updated according to the release *R*64.1.1. Moreover, the updated list of transcription factor binding sites is collected for the same release of genome from SGD. In order to cover more intergenic regions, we consider the whole IGRs that overlap with bound probes and of length more than 30 bp. The mapping of probes on IGRs and the genes regulated by each IGR were obtained from Harbison *et al.* (2004). The sets of all intergenic regions and all probes are denoted by Ω^*I*^ and Ω^*P*^, respectively. For the purpose of our study,two sequence-sets are needed for each ChIP-chip experiment 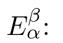:bound sequences and unbound sequences. Therefore, two sequence-sets are defined for each experiment:the bound sequence-set 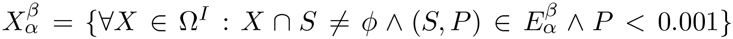 and the unbound sequence-set 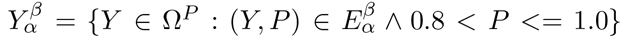. Note that the cardinality of 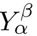 should be five times the cardinality of 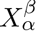 where the probes of the largest p-values are taken first.

To meet the requirements of our study, 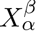 for each ChIP-chip experiment must also meet all the following conditions

1. *β* is a transcription factor protein with a known motif model (PFM),
2. 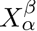 contains only intergenic sequences that regulate verified open reading frames (ORFs),
3. 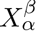 has at least 50 known binding site in SGD, and
4. the cardinality of 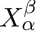 is greater than or equal to 50.

Applying the previous criteria on the 350 ChIP-chip experiments resulted in 38 ChIP-chip datasets corresponding to 22 TFs with known binding motifs. Eventually, one ChIP-chip sequence-set is selected for each TF based on the largest number of *known* binding sites. The final 22 sequence-sets are used to evaluate the performance of our proposed algorithms. Table 3 shows the number of sequences, average sequence length, total number of nucleotides, number of known binding sites and the width of motifs for the bound sequence-sets 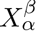. The PWM (log-odds) models of the 22 TF proteins were downloaded from the supplementary materials of MacIsaac *et al.* (2006). However, FSCAN requires PFM (nucleotide frequencies) models in order to calculate K-mer features. A PWM model is converted into a PFM model in three steps. First, the frequencies of the four DNA nucleotides *A,C,G,T* in the intergenic regions set Ω^*I*^ are calculated. Second, PFM entries are generated from PWM scores using the following formula

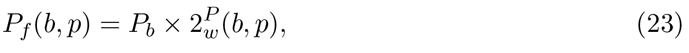

where *P_b_* is the frequency of base *b* in Ω^*I*^. Third, each column *p* of the PFM model is normalized by dividing its elements by Σ_*b*_ *P_f_*(*b,p*). Motif matrices are also trimmed based on the IUPAC consensus sequences Cavener (1987) of their motifs. The head and tail positions that correspond to ‘·’ (any) in the consensus sequences are trimmed. For example, the first and last columns of the motif matrix of AFB1 are removed because the consensus sequence of ABF1 is **rTCAyt.y.ACG.**. This is the strategy used to generate the list of TFBSs in SGD.

**Table 3.**
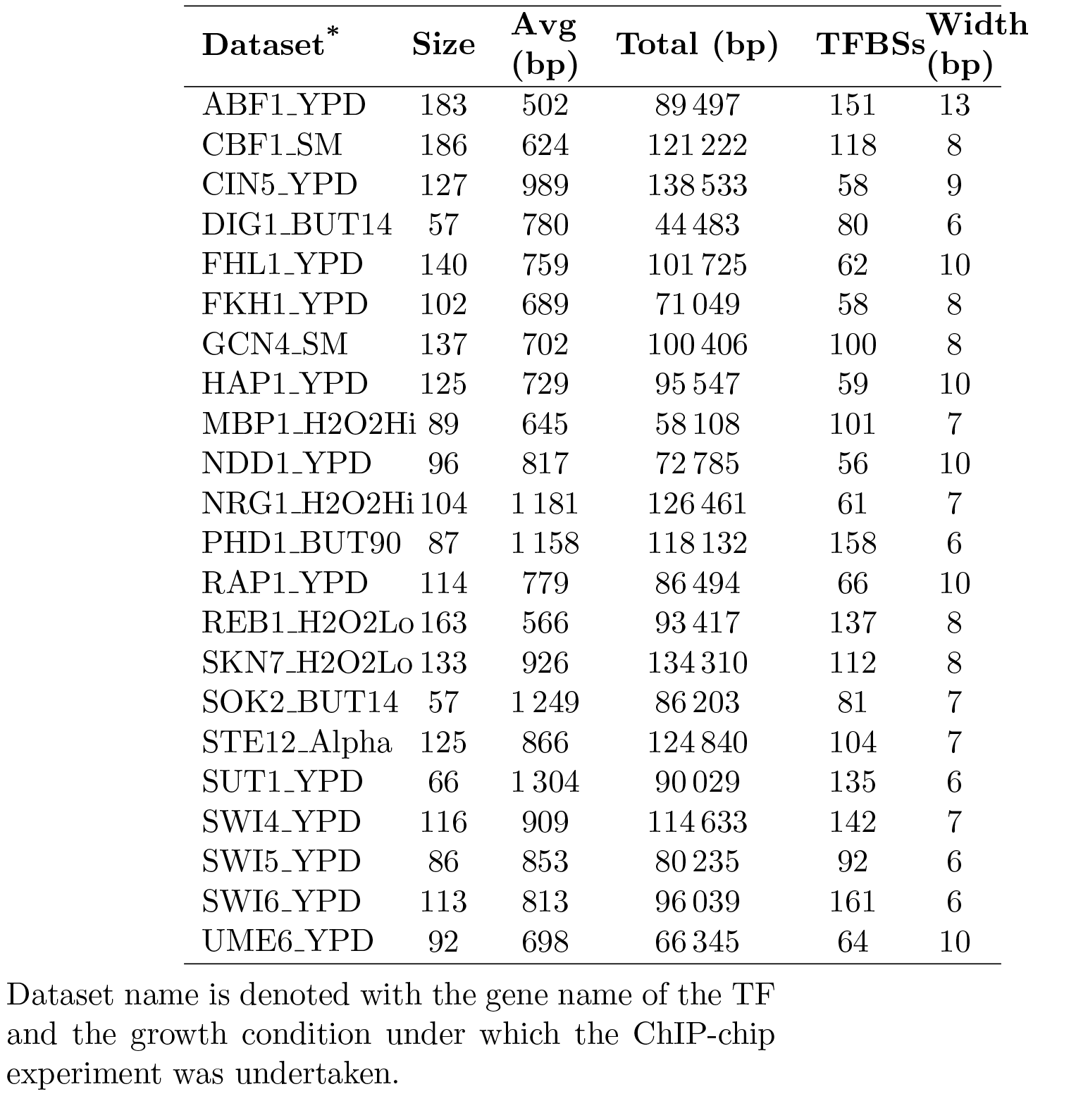
Description of bound sequence-sets 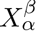 for all datasets.

### 4.2 Evaluation Metrics

Different metrics can be used to evaluate the performance of TFBSs predictors Tompa *et al.* (2005). However, evaluation metrics that can assess the ability of a predictor to discriminate functional K-mers from non-functional K-mers are needed. As a result, we use four performance indexes that are commonly used in machine learning practices to evaluate classification systems. Precision (P) measures the exactness, i.e., the ability of a system to predict binding sites correctly (true positives) while it reduces the number of wrongly predicted binding sites (false positives). Recall (R) measures the completeness,i. e., the ability of a system to predict binding sites correctly while it reduces the number of known binding sites that are wrongly classified (false negatives). To obtain a meaningful measurement of precision and recall, the harmonic mean of both of them is used to measure the effectiveness of classification, called the F1-measure. The performance coefficient (PC) is also used to measure the ability of a system to predict binding sites correctly while it reduces the number of wrongly predicted K-mers. The four evaluation metrics are defined as follows

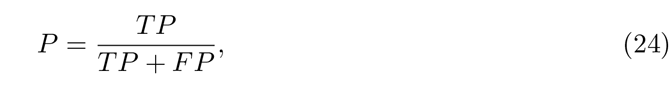

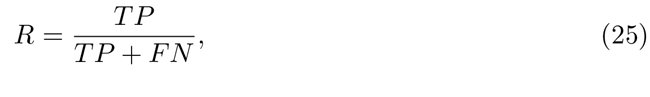

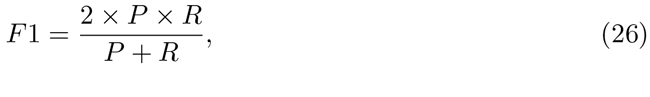

and

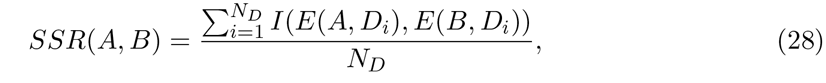

It is worth mentioning that we count a K-mer correctly predicted as a binding site if it exactly matches a known site regardless of the strand. Since the performance of FSCAN is evaluated on a collection of sequence-sets, the average of each measure over all datasets is calculated. To compare the performance of two systems, a system success ratio (SSR) is defined using th following formula

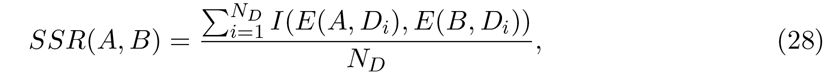

where *E*(*A,D_i_*) is the evaluation metric of system *A* on a sequence-set *D_i_*, *I*(*a,b*) returns 1 iff *a* > = *b*, and N_*D*_ is the total number of datasets.

### 4.3 Results with Comparisons

In order to evaluate the effectiveness and efficiency of our proposed approach, we compare the performance of FSCAN with two popular approaches, MatInspector and MATCH, on the 22 ChIP-chip sequence-sets. Our FSCAN system is run using the default values for all its parameters, i.e., 60% of the motif width positions are selected to generate the DNA-BAS over 100 random trails, 3 membership functions for each of the linguistic variables of the FIS, 0.25 reduction factor in RANDOVER and 3 minority examples are synthesized in SMOTE. On the other hand, we select the evaluation measures at the best performing threshold for each sequence-set when it is tested on MatInspector or MATCH. Table 4 shows the performance measures of FSCAN and the other two approaches. MatInspector and MATCH performs quite similarly in most of the datasets. Table 4 clearly shows that our system FSCAN performs well when the RANDOVER algorithm is used to balance the training data. However, FSCAN has a better performance when the SMOTE algorithm is used to reduce the imbalance between the background K-mers and functional K-mers. From Table 4, it can be seen that our method has higher precision (P) than MatInspector and MACTH in most of the tested sequence-sets (comparing the P columns). In other words, FSCAN has a good ability to reduce the number of wrongly predicted background K-mers (false positives). This ability comes from its strong features that characterize true binding sites as well as the learning ability of ANFIS in generating a well-generalized fuzzy rule base for the classifier fuzzy inference model. Moreover, the recall (R) of FSCAN is better than MatInspector and MATCH when RANDOVER is used to reduce the imbalance effect of training data (comparing the R column of FSCAN-RANDOVER with the R columns MatInspectror and MATCH). When SMOTE is used to handle the imbalance data issue, the performance of FSCAN slightly drops for some datasets. It is worth mentioning that MatInspector and MATCH perform favourably when the scanned motif is highly conserved. For example, the motifs of *DIG1, MBP*1, *SWI*6 and *UME6* are very strong (see Table 9) and consequently MatInspector and MATCH achieves very high recall on the corresponding datasets (see *DIG*1-*BUT14*, *MBP*1_*H*2*O*2*Hi*, *SWI*6_*YPD*, and *U M E*6*_Y_PD* in Table 4). This observation is expected because MatInspector and MATCH use a positional conservation-based similarity metric to scan DNA sequences. On the other hand, FSCAN can retrieve more true binding sites even though the scanning motif is quite weak (see *HAP* 1 and *CIN*5 in Table 9 and *HAP*1_*YPD* and *CIN*5_*JYPD* row in Table 4).

**Table 4:**
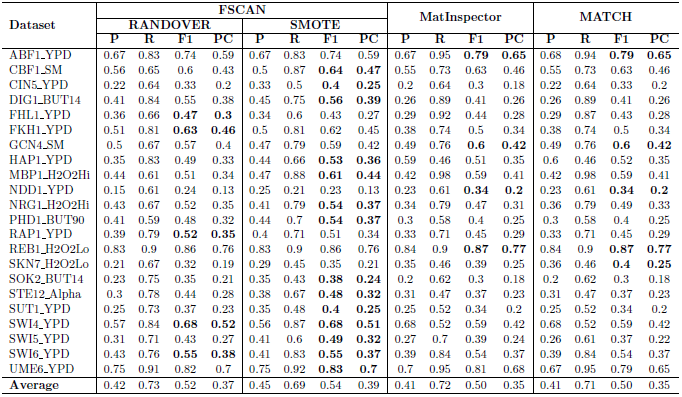
Comparison of FSCAN, MatInspector and MATCH

To recap the performance of our system against MatInspector and MATCH, the F1-measures and performance coefficient (PC) measures are compared. It can be easily seen from Table 4 that FSCAN achieves promising results on 18 datasets and outperforms MatInspector and MATCH in terms of *F*1 and *PC* (see the boldface figures). This means that FSCAN has a very good ability to find true binding sites in the scanned sequences while it reduces the number of wrongly predicted binding sites (false positives) and the number of excluded binding sites (false negatives). Eventually, the success ratio of FSCAN (given in Eq. 28) is calculated against MatInspector and MATCH using the F1-measure as an evaluation metric. Table 5 shows the results of the SSR comparisons. It is obvious that FSCAN achieves better prediction accuracy than MatInspector or MATCH on the majority of the tested sequence-sets. Moreover, the usage of SMOTE gives FSCAN a greater advantage over MatInspector and MATCH. The performance of FSCAN can be further improved if different parameter values are used for each sequence-set. In the next section, further analysis is conducted on the effect of each parameter on the performance of FSCAN.

**Table 5:**
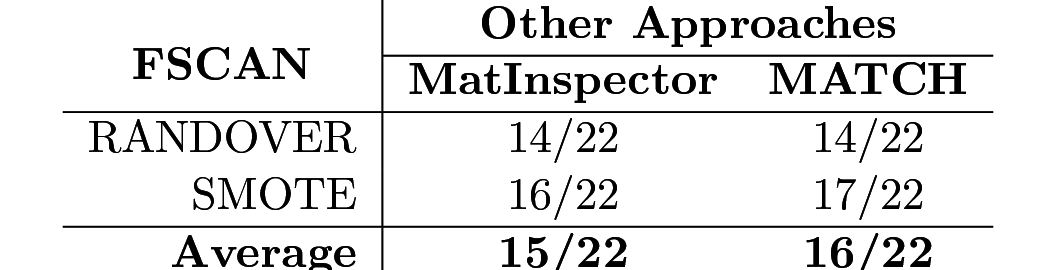
System success ratio (SSR) of FSCAN against Others

### 4.4 Fuzzy Inference of TF DNA-Binding

In this section, the fuzzy inference system (FIS) of one of the tested dataset (DIG1_BUT14) is examined closely in order to acquire some knowledge on how fuzzy rules help predict transcription factor binding sites. We first examine the three membership functions of each fuzzy input variable that are learnt by the ANFIS algorithm. Fig. 4 shows the initial generalized bell-shaped membership functions for each variable that are set using grid partitioning as well as the final membership functions that are used in the FIS. The membership functions of all input variables are initialized similarly so that they span the whole input space. After running the ANFIS algorithm using training and validation datasets, the learning algorithm converges with new membership functions (plotted with solid lines in Fig. 4). It can be noticed that the fuzzy sets for the mean and width of DNA-BAS have quite similar membership functions. However, the small fuzzy set of the mean variable tends to be strict and the large fuzzy set is inclined to be tolerant while the corresponding fuzzy sets of the width variable show the opposite behaviour. The medium fuzzy set for the mean variable is centered around 0.4 while that of the width variable is centred around 0.55. These observations ensure that the mean of the DNA binding affinity signal of TF is smaller than the width of the binding affinity signal in potential binding sites. On the other hand, the sequence conservation variable that represents the phylogenetic footprinting of a K-mer across multiple species shows insignificance for lowly conserved K-mers while K-mers with medium and high conservation scores tend to be more important. This result affirms the fact that functional DNA sequences are highly conserved in related species Blanchette and Tompa (2002).

**Figure 4:**
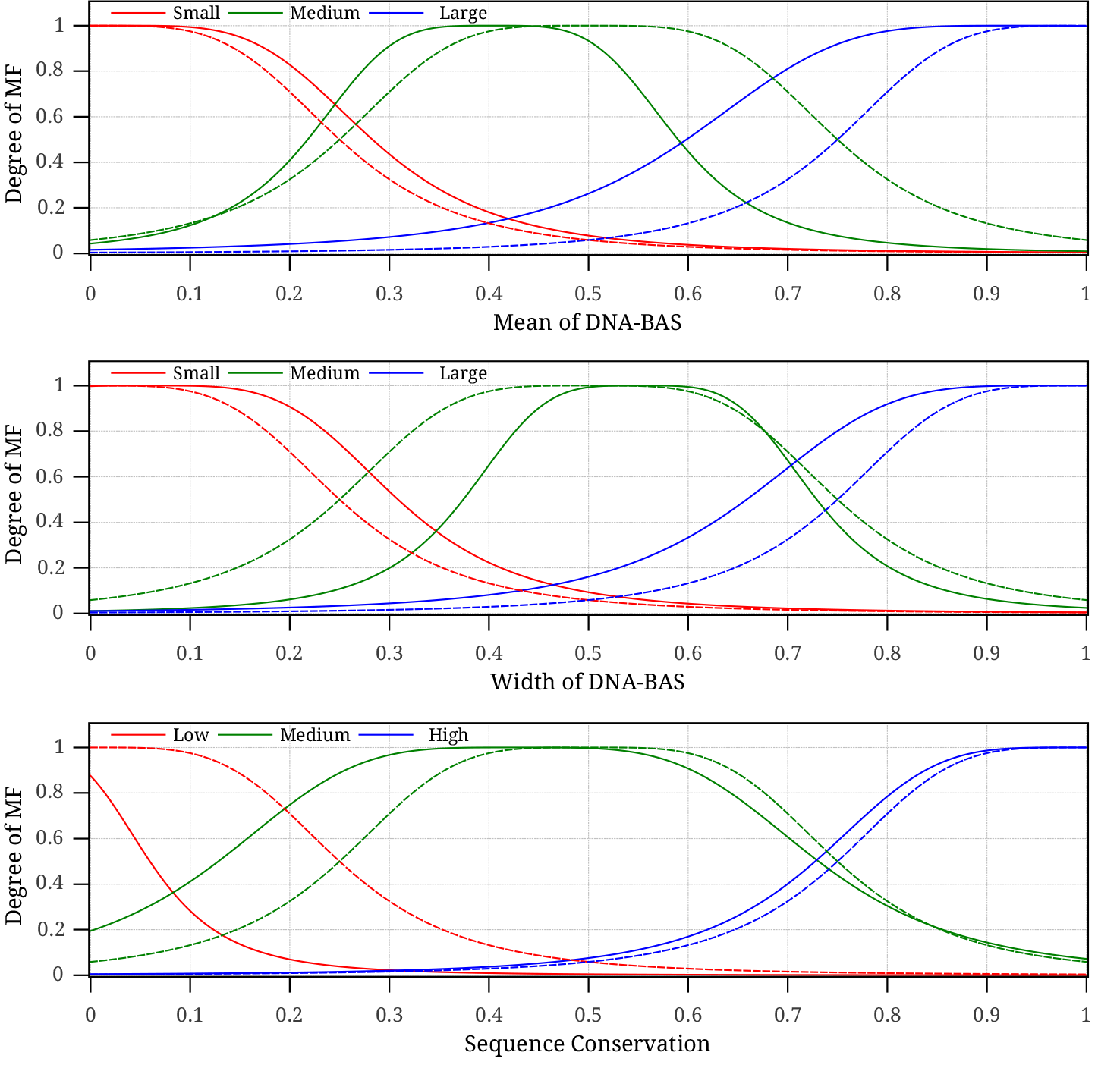
The initial (dashed lines) and final learnt (solid lines) membership functions for the fuzzy input variables of the DIG1_BUT14 dataset.

Next, we show the fuzzy rules that were generated using the ANFIS algorithm. Since grid partitioning is used to initialize the FIS, the number of rules grows exponentially with the number of membership functions and input variables. For simplicity, we use only two membership functions for each fuzzy input variable and a zero-order Sugeno-type inference system. Therefore, each If-Then rule has three parts in the antecedent combined using the fuzzy AND operator and one constant output in the consequent. The eight rules that are generated for DIG1 FIS are presented in Table 6.

**Table 6:**
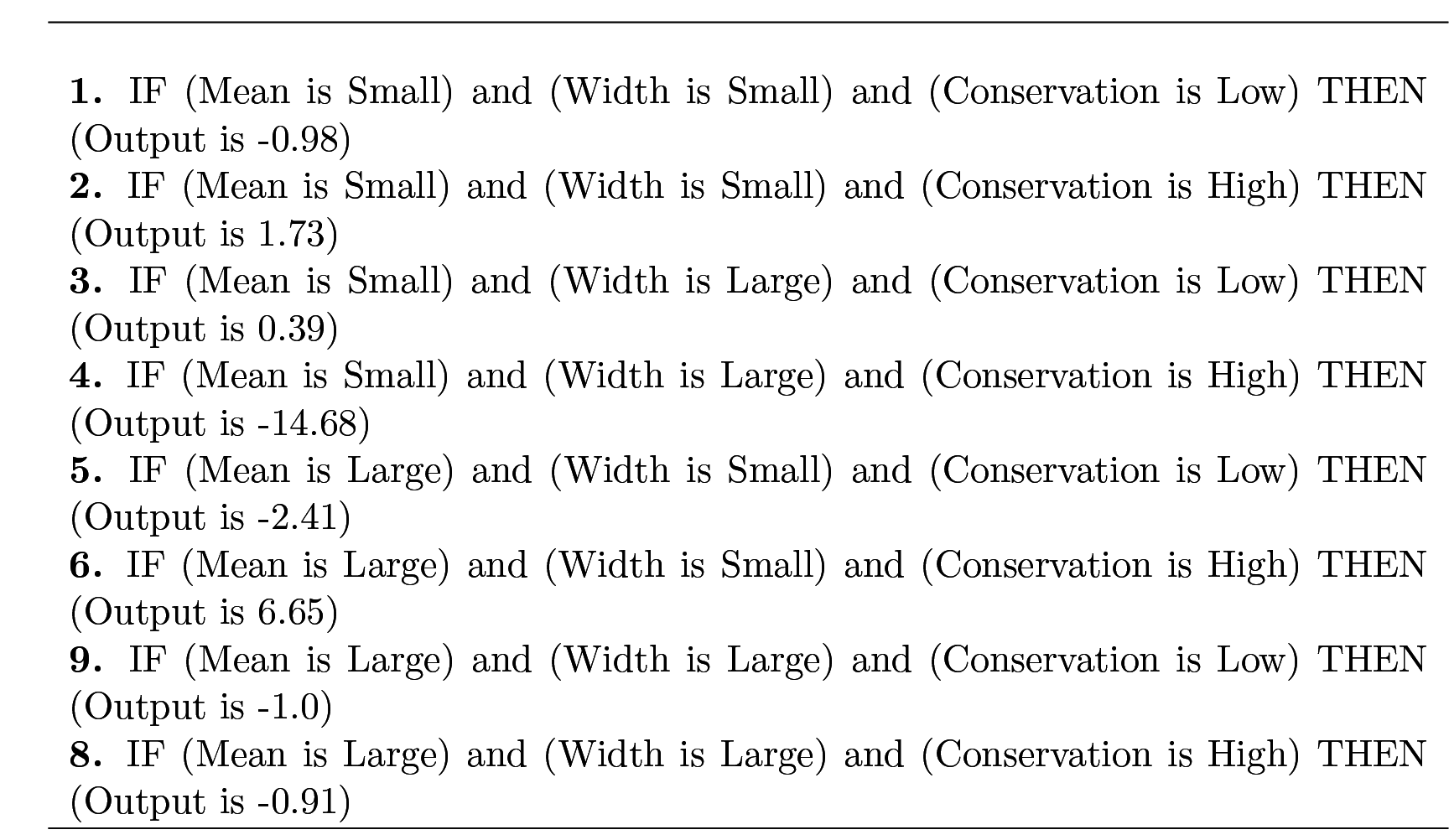
Rules base of DIG1 fuzzy inference system.

When a K-mer modelled by three values is introduced to the FIS, the degree of membership to each fuzzy set of each input variable is computed and then the antecedents of rules are evaluated using the AND fuzzy operator. We use *product* to implement the AND operation and compute the firing strength *w_i_* of the the *i*th rule, i.e., *w_i_* = *μ_m_* × *μ_w_* × *μ_cons_*. The *product* is also used in the implication method in order to scale the constant output levels *z_i_*, i.e., *w_i_* × *z_i_*. The final output of FIS, denoted by *TFBinding*, is the weighted average of all rule scaled outputs

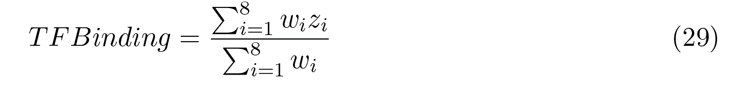

Fig. 5 shows the output surface oi this FIS as the input values vary between [0,1]. Since only positive inference outputs are considered putative binding sites, it can be seen from Fig. 5a that that TF is considered bound to the K-mer only when the width of the DNA-BAS is small or the mean is small given that the sequence conservation is relatively high. We fixed the sequence conservation at 0.5, which is considered fairly high, to produce Fig. 5a. This observation confirms that K-mers with more than 60% of their nucleotide positions similar to the TF motif tend to be putative binding sites. To investigate the impact of the sequence conservation on the TFBS prediction, Fig. 5b shows how the FIS output changes with respect to the mean of DNA-BAS and conservation scores while the width is fixed at 0.4, which is considered quite large for known TFBSs. It can be inferred from this figure that when more than half K-mer nucleotides are dissimilar to the TF motif, the phylogenetic conservation and the mean determine whether the TF binds this K-mer or not. The prediction of TFBS is mainly driven by the second and sixth rule of the rule base in Table 6. Since it is unlikely to have putative binding sites with a large width and small mean, the fourth rule prevents FSCAN from identifying K-mers in this category as TFBSs.

**Figure 5:**
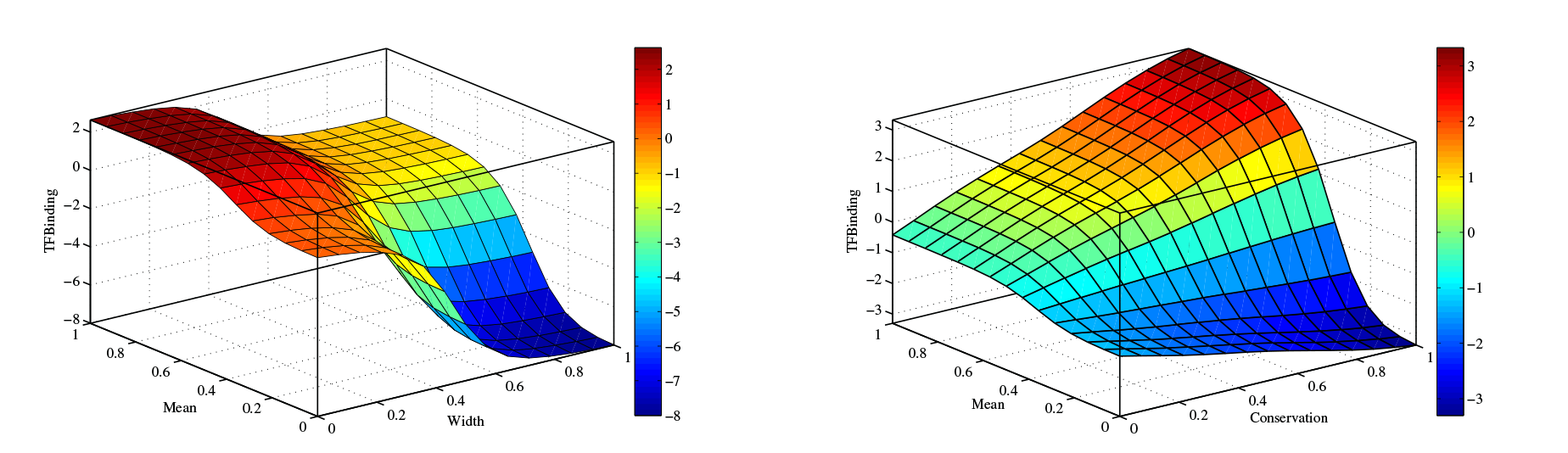
The output surface of FIS when the sequence conservation is fixed at 0.5 and the width of the DNA-BAS is fixed at 0.4, respectively.

The learnt fuzzy rules have given us a good understanding of the TF DNA-binding specificity. To compare the fuzzy rules with the threshold-based rules of MatInspector, we present the crisp rules that are used for DIG1 in Table 7.

**Table 7:**
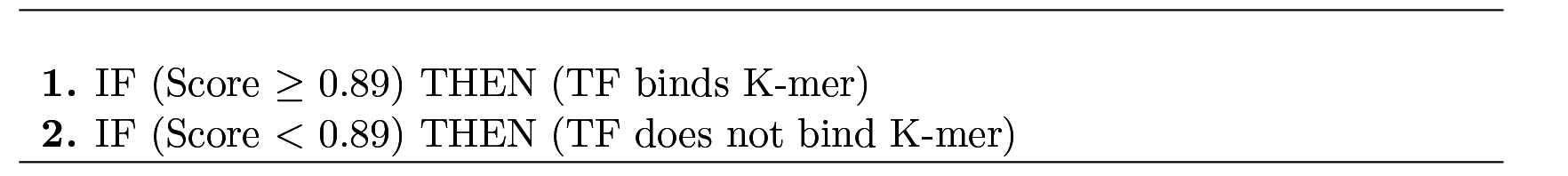
Rules used in MatInspector for DIG1 dataset.

It is obvious from Table 7 that MatInspector is quite sensitive to the threshold value and produces a lot of false positives even though the best threshold is selected.

### 4.5 Robustness Analysis and Discussion

An extensive analysis is performed on FSCAN to investigate the performance of our proposed system in different configurations. The effect of each parameter on the performance of FSCAN is investigated by running FSCAN using different values for each parameter. When the effect of a parameter is analyzed, default values are used for the remaining parameters and SMOTE is used by default to reduce the imbalance ratio in all experiments. F1-measure is used as an evaluation metric in this robustness analysis.

First, the effect of the number of randomly selected positions on performance is inspected. Fig. 6 illustrates how the number of selected positions affect FSCAN’s performance. It can be seen that the performance dramatically changes for some sequence-sets as the number of positions ranges between 10% and 90% of the motif width. For example, the performance of FSCAN on the dataset *CIN*5_*YPD* changes from 0.29 to 0.4 for 10% to 70% of the motif width, respectively. Similarly, F1-measure on *HAP*1_*YPD* ranges between 0.46 and 0.56 for 10% and 50% of the motif width, respectively. On the other hand, FSCAN shows quite robust performance as the number of selected positions change in four sequence-sets (see *FKH*1_*YPD*, *MBP*1_*H*2*O*2*Hi*, *SWI*6-*YPD* and *REB_1H2O2Lo* in Fig. 6). The performance change of FSCAN, over a different number of selected positions, on the rest of the datasets was between 4% and 8%. It can be seen also that FSCAN performs well on most of the datasets when the number of selected positions is less than 60% of the motif width (as shown in a comparison of the black and gray symbols in Fig. 6). Moreover, we calculated the correlation coefficient between the performance changes of FSCAN and the motif widths for all sequence-sets and found it is 0.24. Therefore, there is no strong relationship between the motif width and the performance of FSCAN at a different number of selected positions. Surprisingly, we found that there is a 0.44 correlation between the MISCORE motif scores (*R(M*) in Table 9) and the change in FSCAN performance, as shown in Fig. 6.

**Figure 6:**
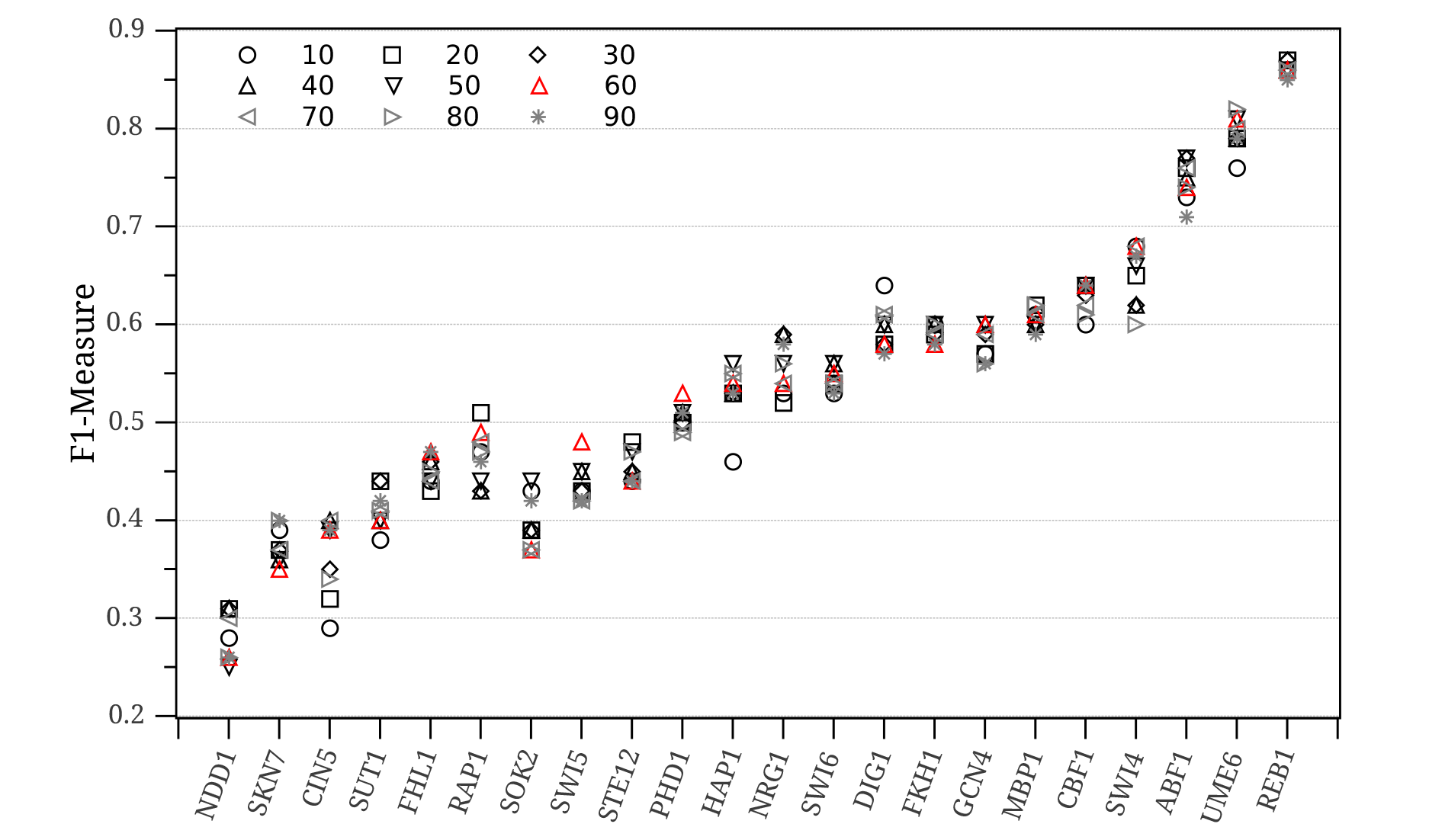
FSCAN performance (F1-Measure) at different a number of selected positions (percentage of the motif width)

Second, the number of random trials that are used to generate the DNA-BAS is investigated. We tried 100, 200, 300, 400 and 500 random trials to generate the affinity signal. Table 8 shows F1-measures for all datasets and for a different number of random trials. It is evident that FSCAN is quite robust with respect to the number of random trials and 100 – 200 trails are sufficient to model the DNA binding affinity for most of the TF proteins. The change in the FSCAN performance over a different number of trials is less than 3% in most cases. Only two TFs show peculiar behavior when the number of random trials becomes greater than 200 (*NRG*1*_H_*2*O*2*Hi* and *RAP*_1*_y_PD* in Table 8).

**Table 8:** FSCAN performance (F1-Measure) at different numbers of random trials R.

| Dataset ID | Number of Random Trials ($R$) | | | | |
| --- | --- | --- | --- | --- | --- |
|  | 100 | 200 | 300 | 400 | 500 |
| **ABF1_YPD** | 0.74 | 0.74 | 0.74 | 0.73 | 0.73 |
| **CBF1_SM** | 0.64 | 0.64 | 0.63 | 0.64 | 0.64 |
| **CIN5_YPD** | 0.39 | 0.39 | 0.38 | 0.37 | 0.36 |
| **DIG1_BUT14** | 0.58 | 0.58 | 0.61 | 0.61 | 0.61 |
| **FHL1_YPD** | 0.47 | 0.47 | 0.47 | 0.46 | 0.48 |
| **FKH1_YPD** | 0.58 | 0.6 | 0.59 | 0.6 | 0.6 |
| **GCN4_SM** | 0.6 | 0.6 | 0.6 | 0.6 | 0.6 |
| **HAP1_YPD** | 0.54 | 0.55 | 0.54 | 0.54 | 0.53 |
| **MBP1_H2O2Hi** | 0.61 | 0.61 | 0.6 | 0.62 | 0.61 |
| **NDD1_YPD** | 0.26 | 0.25 | 0.27 | 0.25 | 0.26 |
| **NRG1_H2O2Hi** | 0.54 | 0.57 | 0.53 | 0.56 | 0.55 |
| **PHD1_BUT90** | 0.53 | 0.53 | 0.53 | 0.52 | 0.53 |
| **RAP1_YPD** | 0.49 | 0.43 | 0.43 | 0.45 | 0.44 |
| **REB1_H2O2Lo** | 0.86 | 0.86 | 0.85 | 0.87 | 0.86 |
| **SKN7_H2O2Lo** | 0.35 | 0.36 | 0.35 | 0.35 | 0.36 |
| **SOK2_BUT14** | 0.37 | 0.39 | 0.37 | 0.39 | 0.38 |
| **STE12_Alpha** | 0.44 | 0.46 | 0.45 | 0.44 | 0.45 |
| **SUT1_YPD** | 0.4 | 0.41 | 0.42 | 0.41 | 0.42 |
| **SWI4_YPD** | 0.68 | 0.67 | 0.65 | 0.67 | 0.66 |
| **SWI5_YPD** | 0.48 | 0.47 | 0.5 | 0.5 | 0.47 |
| **SWI6_YPD** | 0.55 | 0.56 | 0.55 | 0.54 | 0.54 |
| **UME6_YPD** | 0.81 | 0.81 | 0.8 | 0.81 | 0.81 |

Third, the performance of FSCAN is analyzed at a different number of membership functions for the input linguistic variables of the FIS that is learnt by ANFIS. Fig. 7 depicts the F1-measures of FSCAN at 2 (small, large),3 (small, medium, large) and 4 (small, medium, large, very large) membership functions for each testing ChIP-chip sequence-set. Surprisingly, 12 TF proteins of the 22 studied proteins show a good performance for FSCAN when only two membership functions are used to generate the fuzzy rule base. These datasets are: *ABF*1_*YPD*, *CIN*5_*YPD*, *FKH*1_*YPD*, *GCN*4_*SM*, *NDD*1_*YPD*, *NRG*1_*H*2*O*2*Hi*, *RAP*1_*YPD*, *REB*1_*H*2*O*2*Lo*, *SKN*7_*H*2*O*2*Lo*, *SOK*2_*BUT*14, *STE*12_*Alpha*, and *UME*6_*YPD* in Fig. 7. This can be attributed to the discrimination power of the K-mer features that are used as inputs to the fuzzy inference system. However, FSCAN performs favourably on nine datasets when three membership functions are used (see *CBF*1_*SM*, *FHL*1_*YPD*, *HAP*1_*YPD*, *MBP*1_*H*2*O*2*Hi*, *PHD*1_*BUT*90, *SUT*1_*YPD*, *SWI*4_*YPD*, *SWI*5_*YPD*, and *SWI*6_*YPD* in Fig. 7). Only one sequence-set significantly benefits from increasing the number of membership functions for the system linguistic variables (as shown in comparison of the blue bar with the green and red bars of *DIG*1 in Fig.7).The correlation coefficient between the best number of membership functions for each sequence-set and its motif width is –0.5. In other words, short motifs seem to require more membership functions in their FISs in order to capture the true binding sites. This is the case in *DIG*1_*BUT*14.

**Figure 7:**
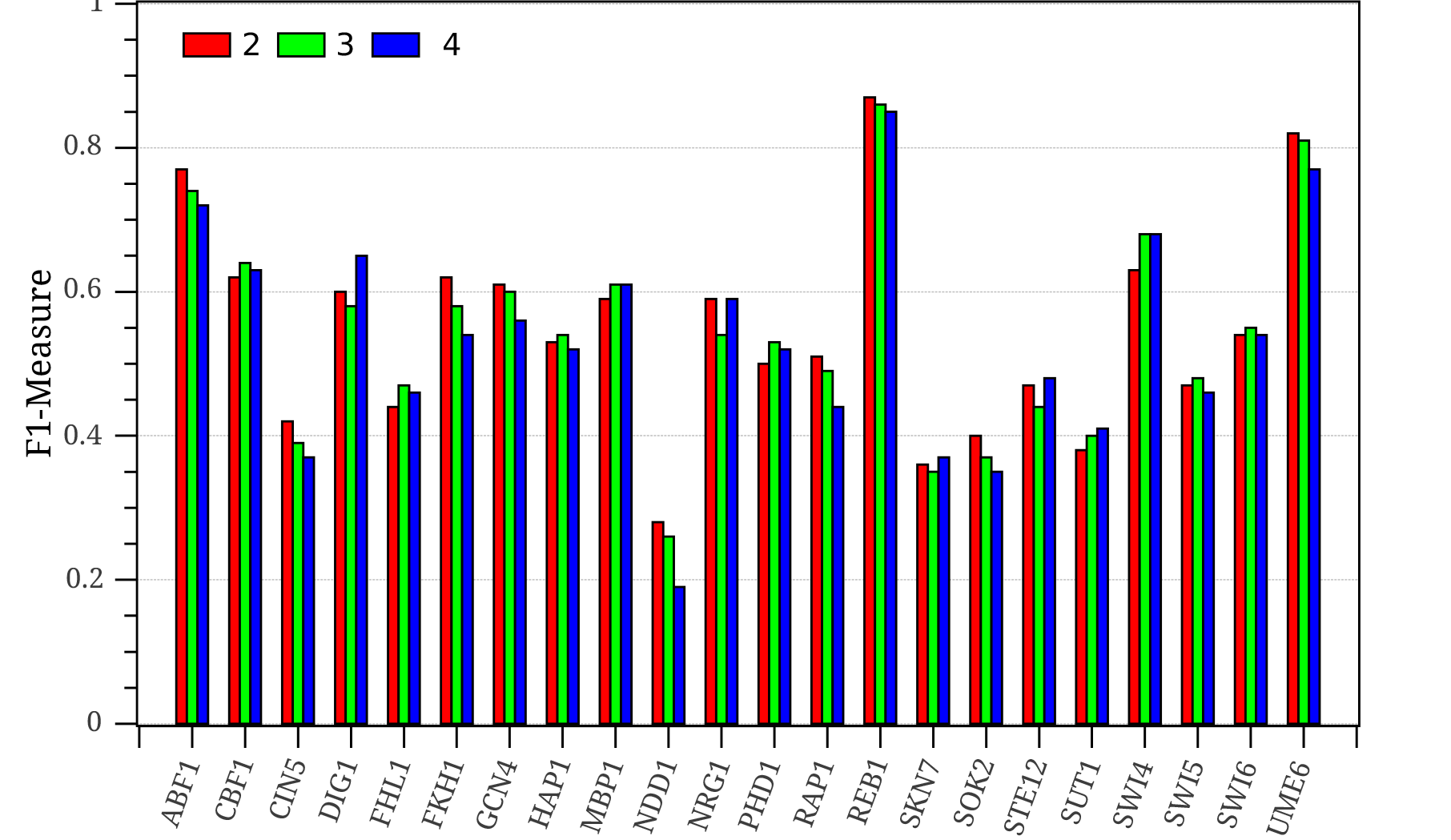
FSCAN performance (F1-Measure) at 2, 3, and 4 fuzzy membership functions for each input.

Finally, the effect of the imbalance ratio on FSCAN performance is examined. Fig. 8 and Fig. 9 show F1-measures of FSCAN at different values for the reduction factor *β* of RANDOVER (Algorithm 2) and at different percentages of synthesized examples in SMOTE, respectively. In both figures, the dashed line represents the performance of FSCAN when the imbalance reduction functionality is disabled, i.e., the ANFIS learns the fuzzy rule base using imbalanced training datasets. First, the performance of FSCAN when the RANDOVER algorithm is used to reduce the imbalance between background K-mers and binding sites is analyzed. It is obvious that FSCAN, with RANDOVER regardless of the reduction factor value, performs better than FSCAN without the imbalance data reduction algorithm for most of the datasets (as shown in comparison of the dashed line with other symbols in Fig. 8). In *HAP* 1 and *DIG1* datasets, however, the performance of FSCAN degrades below the dashed line if the reduction factor *β* exceeds 0.25. From Fig. 8, it can also be observed that 0 < *β* ≤ 0.25 is sufficient for FSCAN to perform well on most of the sequence-sets as soon as the imbalance ratio between the background K-mers and binding sites is larger than 5 (as shown in BG/BS in Table 2 and the black and red symbols in Fig.8).Two datasets, however, require high a reduction factor (0.85) for FSCAN to produce good performance (see *MBP*1 and *CBF*1 in Fig. 8). The imbalance ratio of these two datasets is quite low (see Table 2) and more minority examples are needed to make the datasets well-balanced for the ANFIS learning algorithm.

**Figure 8:**
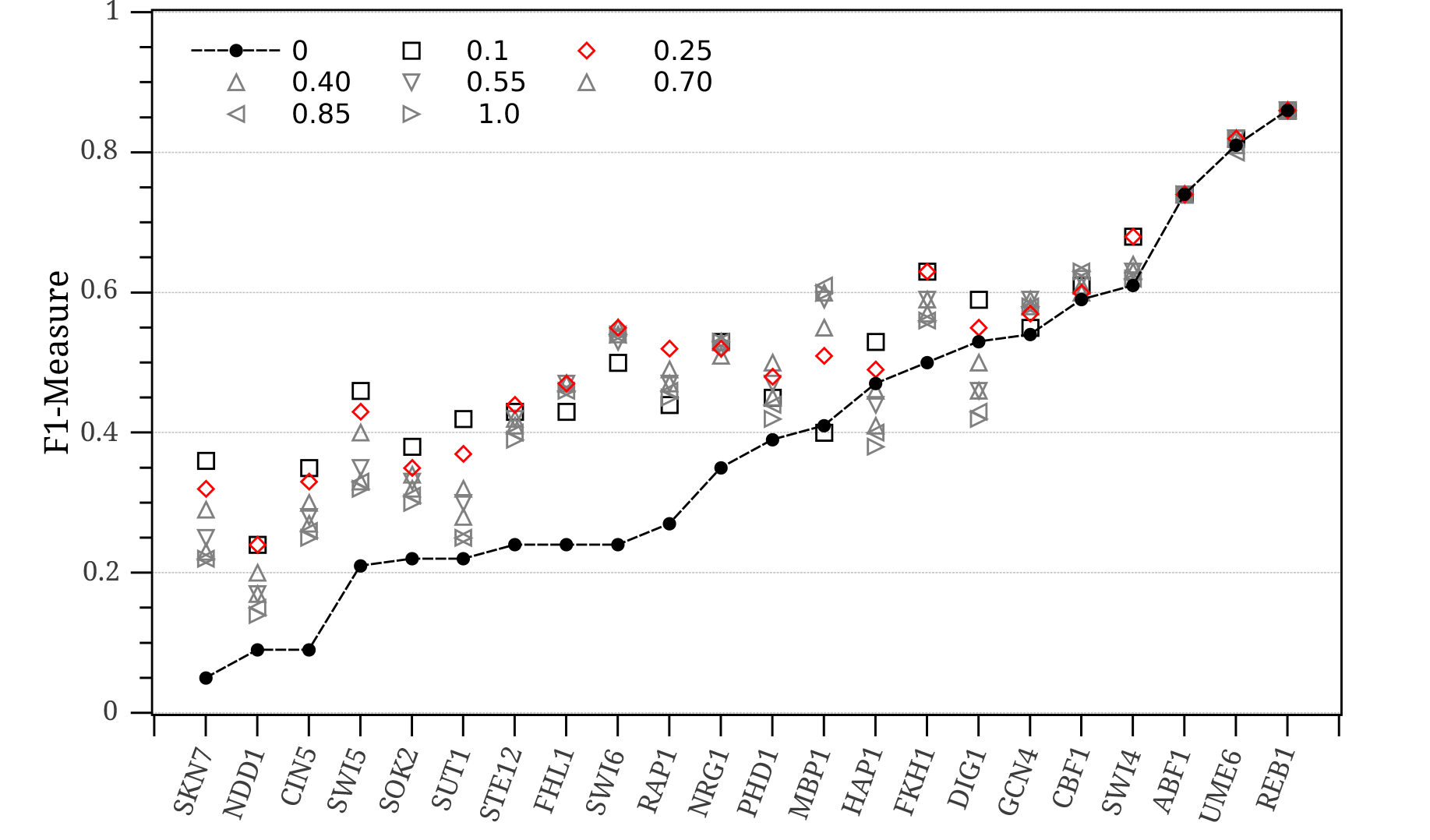
FSCAN performance (F1-Measure) with different values for the reduction factor *P* in th RANDOVER algorithm.

**Figure 9:**
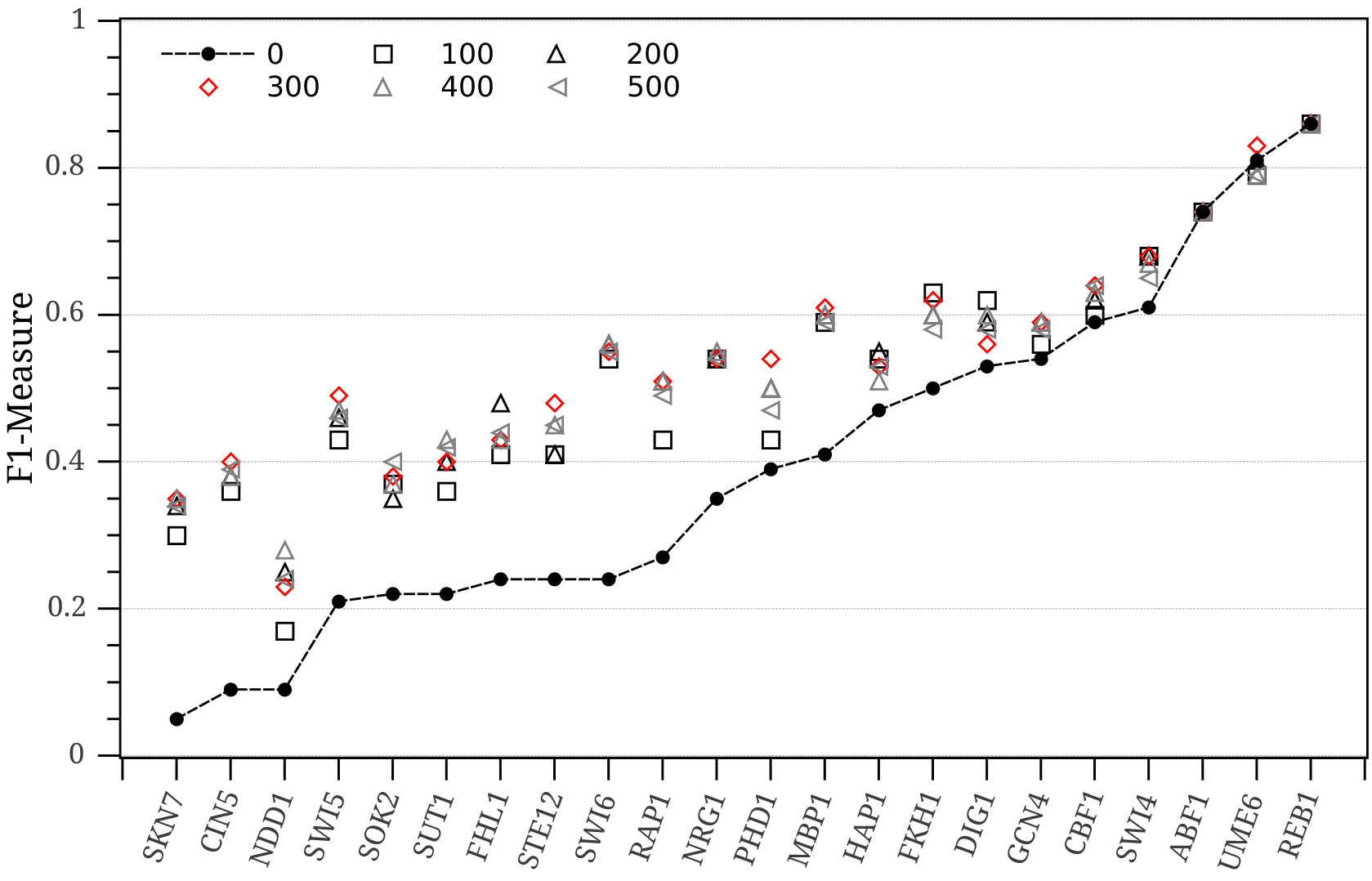
FSCAN performance (F1-Measure) at different percentages of synthesized examples in the SMOTE algorithm

Second, the performance of FSCAN when SMOTE is used for imbalance reduction is studied. SMOTE is tested on different percentages of synthesized examples, i.e., the number of the newly generated examples is a percentage of the number of examples in the minority class. It is clear from Fig. 9 that 100-300% are suitable for FSCAN to perform well on most of the datasets. However, three sequence-sets show a better performance when 400% of the minority class examples are generated (as shown in a comparison of the symbols above *NDD*1, *SUT*1 and *NRG*1 in Fig. 9). Moreover, SMOTE always improves the performance of FSCAN regardless of the synthetic percentage (as shown in a comparison of the dashed line with the other symbols in Fig. 9). When the imbalance ratio is close to 1, the number of synthetic examples should be considered carefully as in the case of *UME*6.

## 5 Conclusion

Identifying transcription factor binding sites (TFBSs) has been a challenging task in computational biology and bioinformatics for a long time. The main hurdle of this research is setting a cut-off threshold that classifies K-mers into functional (TFBS) or non-functional sequences (background). Moreover, most of the existing approaches are solely based on the primary DNA sequence of the binding sites while some approaches try to use prior knowledge of their locations in the studied genome. However, these approaches lack the ability to learn from existing known binding sites.

In this paper, our proposed system, called FSCAN, overcomes these limitations by using a fuzzy inference system (FIS) based approach. The fuzzy rule base and the membership functions of the linguistic variables in FSCAN are learnt from a training dataset using the ANFIS algorithm. The training dataset is created from a set of sequences that have known binding sites. We proposed a novel approach to model the K-mers that are extracted from the sequence-set and convert them into features/label pairs. These features included the mean and width of the DNA-BAS which represents the binding affinity of different combinations of DNA bases to their corresponding positions in the motif PFM. Another prior knowledge feature based on the phastCons scores was used to represent the phylogenetic footprinting of putative binding sites across species. Moreover, we developed an adaptive ellipsoidal filtering (AEF) algorithm that learned a filter model from the data distribution of each TF separately unlike other existing approaches that use a cutoff threshold. AEF helps remove background K-mers effectively while it preserves true binding sites.

Our proposed approach is threshold-free and can integrate more prior knowledge to improve prediction accuracies. Obviously, the performance of FSCAN boosts as more true binding sites become available. The performance of FSCAN was tested on 22 sequence-sets extracted from genome-wide ChIP-chip experiments on the Saccharomyces Cerevisiae genome. The testing results showed superior performance for FSCAN against other traditional approaches (MatInspector Quandt *et al.* (1995) and MATCH Kel *et al.* (2003)). A detailed robustness analysis was conducted on FSCAN to examine the effect of each parameter on its performance. It was evident from the results that some of these parameters, e.g., number of selected positions and imbalance ratio, greatly affected the performance of FSCAN and should be deliberately set for some sequence-sets. On the other hand, the number of random trials in the DNA-BAS and the number of membership functions in the input linguistic variables of FIS did not have an influential effect on FSCAN performance and can be fixed.

In a nutshell, this study ensures that fuzzy inference systems can be very effective in identifying TFBSs in a set of DNA sequences that are bound by a specific TF protein in a ChIP-chip experiment. It also encourages the usage of learning-based models to characterize TFBSs rather than using traditional computational approaches. Moreover, the proposed approach can be easily applied to scan DNA sequences from any genomic region in order to locate putative binding sites. In this case, the over-representation term Δ in Eq. (10) can be set to 1 and there is no need to have the unbound sequence-set. In addition to that, our proposed solution can be extended to other types of protein-DNA interaction data, e.g., ChIP-seq. Future work will include more biological and computational features to model the TFBSs and more TF proteins from different genomes will be tested.

## A Sequence Logos for the Studied TF Motifs

The motif logos of the TF proteins used in this study are presented in Table 9.

**Table 9:**
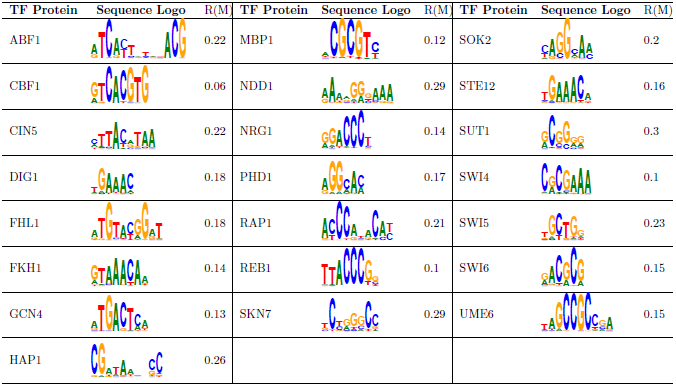
Sequence logos of the investigated TF proteins and their corresponding MISCORE-based motif scores (MMSs).

| TF Protein | Sequence Logo | R(M) | TF Protein | Sequence Logo | R(M) | TF Protein | Sequence Logo | R(M) |
| --- | --- | --- | --- | --- | --- | --- | --- | --- |
| ABF1 |  | 0.22 | MBP1 |  | 0.12 | SOK2 |  | 0.2 |
| CBF1 |  | 0.06 | NDD1 |  | 0.29 | STE12 |  | 0.16 |
| CIN5 |  | 0.22 | NRG1 |  | 0.14 | SUT1 |  | 0.3 |
| DIG1 |  | 0.18 | PHD1 |  | 0.17 | SWI4 |  | 0.1 |
| FHL1 |  | 0.18 | RAP1 |  | 0.21 | SWI5 |  | 0.23 |
| FKH1 |  | 0.14 | REB1 |  | 0.1 | SWI6 |  | 0.15 |
| GCN4 |  | 0.13 | SKN7 |  | 0.29 | UME6 |  | 0.15 |
| HAP1 |  | 0.26 |  |  |  |  |  |  |

## Acknowledgment

M. A. was receiveing a PhD scholarship from La Trobe University and a Top-Up Ph.D. scholarship from Victorian Life Sciences Computation Initiative (VLSCI) during the development of this research.

